# High-density culture of bovine embryonic stem cell–derived mesenchymal cells on edible scaffolds for structured cultivated meat

**DOI:** 10.64898/2026.04.23.720345

**Authors:** Madeleine Carter, Tim WGM Spitters, Sarah Ho, Sophia Webb, Niamh Hyland, Patrick Joseph Mee, Simona Fehlmann, Deepika Rajesh

## Abstract

Developing structured cultivated meat requires integrated solutions that combine scalable cell sources with edible, foodgrade materials capable of supporting highdensity growth and differentiation. Here, we evaluate bovine mesenchymal stem cells derived from embryonic stem cells (ESCderived iMSCs) as a scalable adipogenic cell source and develop an integrated workflow combining these cells with edible plantbased scaffolds for structured biomass generation. Cell identity and functionality were assessed using transcriptomic, morphological, gene expression, flow cytometric, and adipogenic differentiation analyses, in both adherent and suspension culture systems. In parallel, lentil, pea, and soy-based scaffold formulations were screened for cell attachment, proliferation, and biomass accumulation. Soybased scaffolds supported uniform cell distribution and robust growth and outperformed lentil-based scaffolds. Under dynamic culture conditions, bovine iMSCs cultured on soy-based scaffolds achieved highdensity growth, showing biomass accumulation (cell wet weight/scaffold wet weight) reached an average cell wet weight to scaffold wet weight ratio of 15% within three days. Cultures demonstrated active glucose metabolism and retained adipogenic differentiation capacity, confirmed by lipid accumulation and positive oil red O staining. These findings demonstrate an integrated cell–scaffold platform for rapid threedimensional biomass generation. This approach supports the development of a cell culture strategy for structured cultivated meat by combining defined cell sources with foodgrade scaffold technologies to improve scalability, structure, and nutritional relevance.

**Highlights:**

- Bovine ESC-derived iMSCs enable scalable adipogenic cell production
- Edible soy-based scaffolds support 3D attachment and biomass accumulation
- Dynamic culture achieved ∼15% cell wet weight fraction within 3 days
- iMSCs retained adipogenic differentiation capacity on edible scaffolds
- Integrated cell–scaffold culture supports structured cultivated meat prototypes

## Introduction

Cultivated meat manufacturing requires cell sources that can be expanded reproducibly, directed toward relevant tissue phenotypes, and incorporated into edible three-dimensional structures. Cell sourcing is a core consideration across the cultivated meat production workflow, influencing achievable scale, differentiation outcomes, media requirements, and compatibility with downstream structuring approaches (Reiss et al., 2021). In parallel, the sensory and nutritional characteristics of cultivated products—particularly texture, color, flavor, and lipid content—are key determinants of product quality and consumer relevance (Fraeye et al., 2020; Lew et al., 2024).

Current approaches to cultivated meat commonly employ both primary animal cell sources and research model systems. For example, bovine muscle-derived fibro-adipogenic progenitor cells have been shown to support extensive proliferation and robust adipogenic differentiation, including within three-dimensional edible scaffold formats (Dohmen et al., 2022). In method development for structured tissues, murine cell lines such as C2C12 myoblasts and 3T3-L1 preadipocytes are frequently used to establish scaffold designs and fabrication strategies, including stacked muscle-and adipose-like constructs on edible protein-based materials (Li et al., 2022; Shahin-Shamsabadi & Selvaganapathy, 2022; Yao et al., 2024). These model systems are valuable for rapid prototyping of culture and structuring methods; however, translation to scalable livestock-derived cell platforms suitable for food production remains an active area of development (Reiss et al., 2021; Fish et al., 2020).

Mesenchymal stem or stromal cells (MSCs) represent a practical intermediate cell type for cultivated meat applications because they can be expanded *in vitro* and possess mesodermal lineage potential, including adipogenic differentiation. Bovine MSCs isolated from bone marrow have been shown to exhibit multilineage differentiation capacity, including adipogenic commitment under defined culture conditions, supporting their relevance as a livestock-derived cell source (Bosnakovski et al., 2005; Okamura et al., 2018; Hill et al., 2019). Adipose tissue is particularly relevant for cultivated meat products because fat contributes substantially to flavor, texture, and overall organoleptic perception, and recent studies demonstrate that cultivated fat can generate volatile and sensory profiles that overlap with those of conventional animal fat (Fraeye et al., 2020; Lew et al., 2024).

A second major challenge for cultivated meat is structurization; the formation of edible, tissue-like architectures that retain biomass while enabling nutrient transfer and handling during culture. In other biological contexts, self-assembly approaches have demonstrated that multicellular progenitor populations can form higher-order organoid-like structures under defined conditions (Ebner-Peking et al., 2021; Lewis-Israeli et al., 2021). However, producing macroscale structured biomass for food applications typically requires supporting biomaterials that provide mechanical integrity, facilitate cell attachment, and are compatible with manufacturing constraints.

Product-relevant scaffolds for cultivated meat must be edible, scalable, and permissive for cell adhesion, proliferation, and differentiation. Reviews of cultivated meat scaffolding emphasize the importance of food-grade materials and fabrication strategies that avoid scaffold removal and reduce downstream processing complexity (Moslemy et al., 2023). Recent work using decellularized plant-derived scaffolds further demonstrates that edible architectures can support muscle and adipose cell attachment and differentiation and yield prototypes with measurable textural properties (Ben-Arye et al., 2020; Murugan et al., 2024). From a bioprocess perspective, porous edible scaffolds can intensify culture by increasing available surface area for cell adhesion and supporting high biomass accumulation, aligning scaffold design with scalable culture formats (Hanga et al., 2026).

In this study, we evaluate an integrated workflow combining bovine embryonic stem cell-derived mesenchymal stem cells (ESC-derived iMSCs) with edible soybased scaffolds for threedimensional, highdensity culture relevant to structured cultivated meat production (Figure 1). Cell identity, adipogenic competence, scaffold compatibility, and shortterm highdensity culture performance are assessed across a staged experimental framework encompassing cell characterization, scaffold screening, and dynamic culture evaluation.

**Figure 1.**
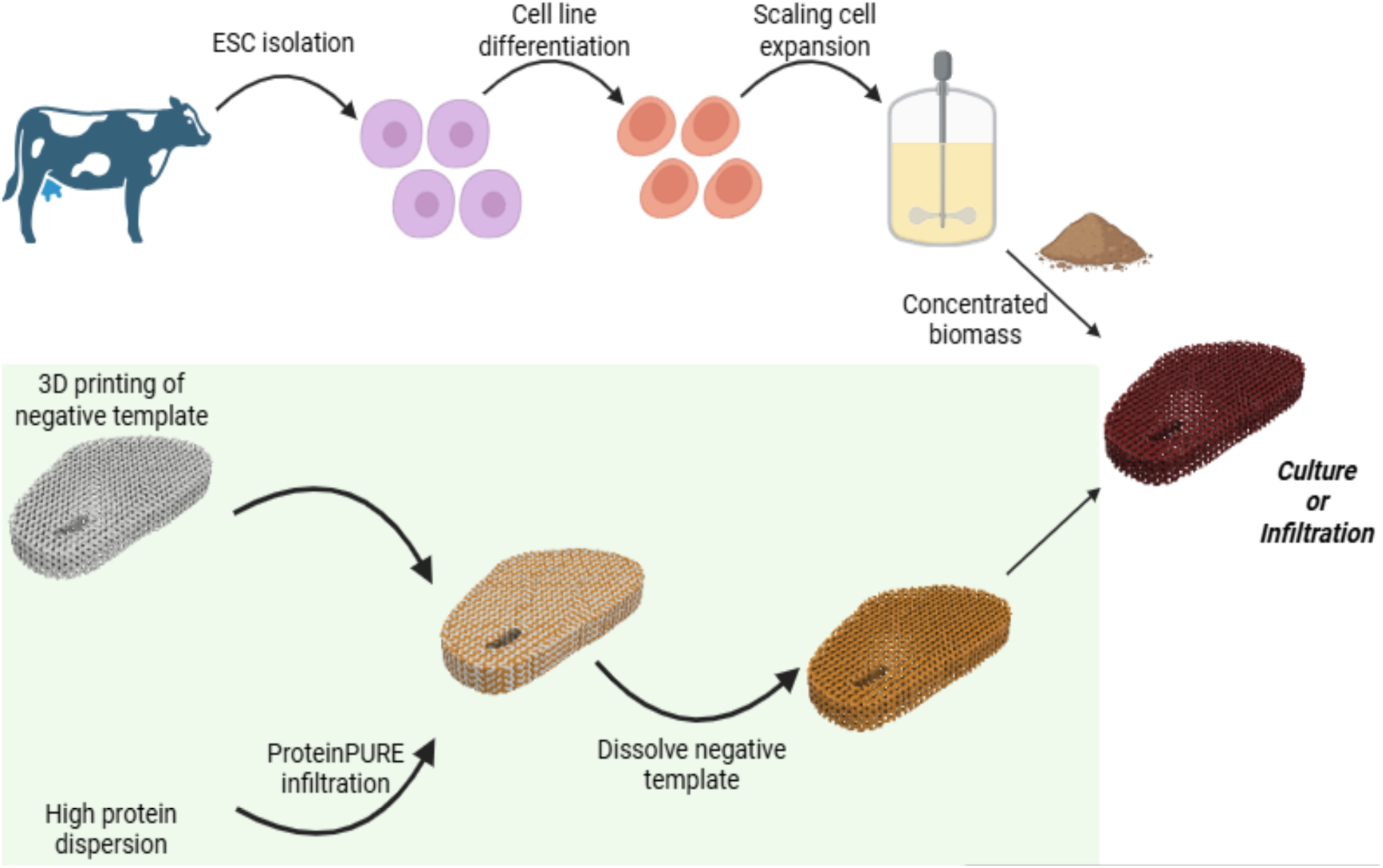
Integrated cell derivation and scaffold-based workflow for three-dimensional biomass generation. Schematic overview of the experimental strategy used to generate structured three-dimensional biomass for cultivated meat applications. Pluripotent bovine embryonic stem cells (ESCs) are isolated, expanded, and differentiated through defined lineage stages to generate mesenchymal stem cells, which are subsequently expanded for biomass production. In parallel, edible plant-protein scaffolds are fabricated using additive manufacturing of a sacrificial template followed by protein infiltration to create defined porous structures suitable for cell culture. The two components are combined by culturing or seeding cells onto the edible scaffolds, enabling three-dimensional cell attachment, expansion, and downstream differentiation within a food-grade matrix.

**Figure 2.**
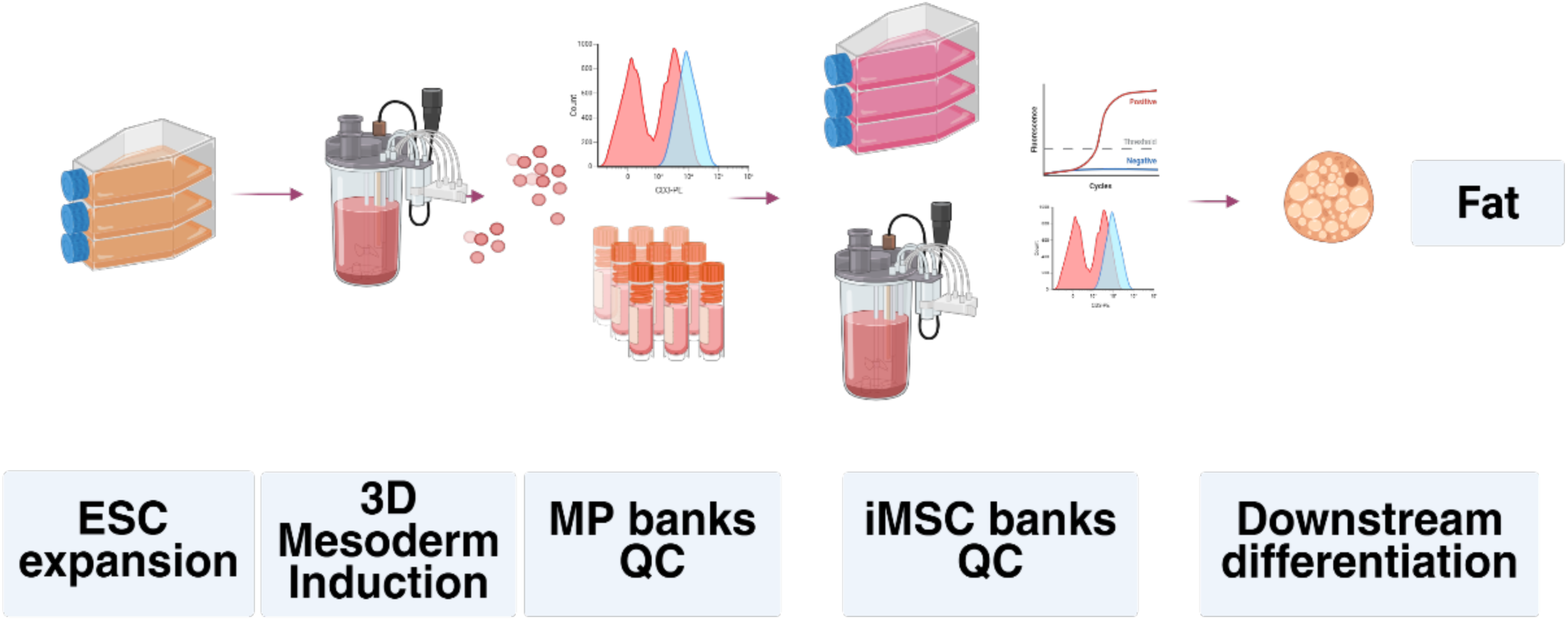
Derivation and phenotypic characterisation of bovine embryonic stem cell–derived mesenchymal stem cells. Pluripotent bovine embryonic stem cells (ESCs) were expanded and induced toward mesodermal fate under three-dimensional (3D) culture conditions, generating mesodermal progenitor (MP) populations that were subsequently expanded and cryopreserved as MP banks. Controlled lineage specification from MP stages yielded induced mesenchymal stem cells (iMSCs), which were further expanded to establish iMSC cell banks following quality control (QC) assessment. Molecular and phenotypic characterisation of banked iMSCs was performed using RT-qPCR and flow cytometry to confirm acquisition of mesenchymal identity. Functional qualification of iMSC banks was achieved by demonstrating the capacity for downstream adipogenic differentiation, supporting their suitability as a renewable mesenchymal cell source for scaffold-based culture applications.

**Figure 3.**
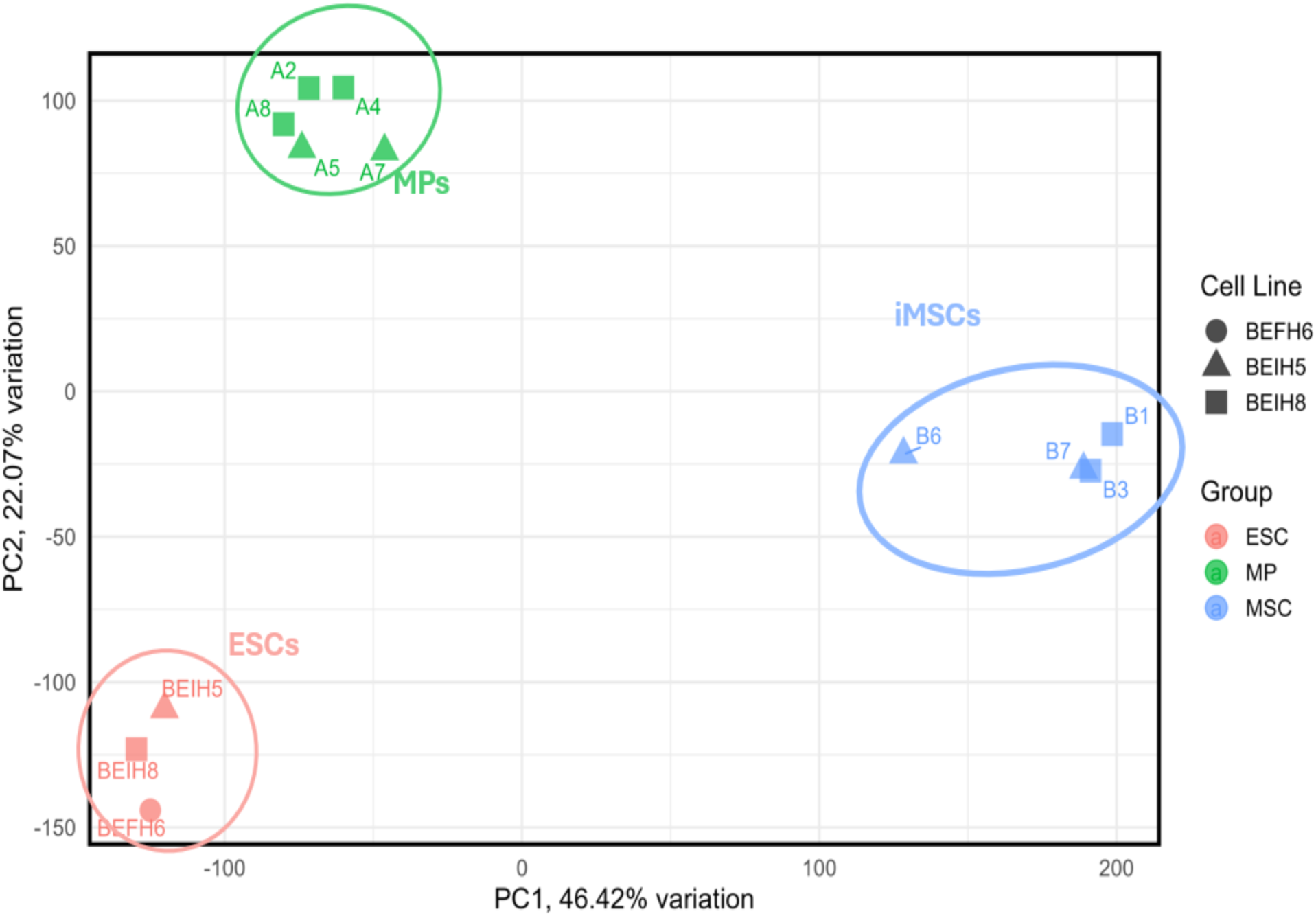
Transcriptomic profiling of bovine embryonic stem cell–derived mesenchymal stem cell differentiation. Transcriptomic profiling of bovine embryonic stem cells (ESCs), mesodermal progenitors (MPs), and induced mesenchymal stem cells (iMSCs) was performed by RNA sequencing and visualised using principal component analysis (PCA). The analysis shows distinct clustering of samples according to differentiation stage, supporting transcriptional divergence during lineage progression from pluripotent ESCs through MP intermediates to iMSCs.

## Materials and Methods

### Cell proliferation assays

Cell number and cell wet weight (ww) were determined using standard curves correlating MTT absorbance with both cell number and biomass. For the 40 min MTT incubation (Figure 5), cell wet weight was linearly related to cell number (cell wet weight = 2.7369 × 10⁻⁹ × number of cells), while exponential relationships were established between MTT absorbance and both cell number (number of cells = 5.5192 × 10⁵ × exp(4.1167 × MTT absorbance)) and cell wet weight (cell wet weight = 5.0319 × 10⁻³ × exp(1.5471 × MTT absorbance)). For the 60 min MTT incubation (Figure 4), similar relationships were obtained, with cell wet weight linearly correlated to cell number (cell wet weight = 2.6138 × 10⁻⁹ × number of cells), and exponential correlations observed for cell number (number of cells = 2.7918 × 10⁵ × exp(3.8473 × MTT absorbance)) and cell wet weight (cell wet weight = 9.8314 × 10⁻⁴ × exp(3.3955 × MTT absorbance)).

**Figure 4.**
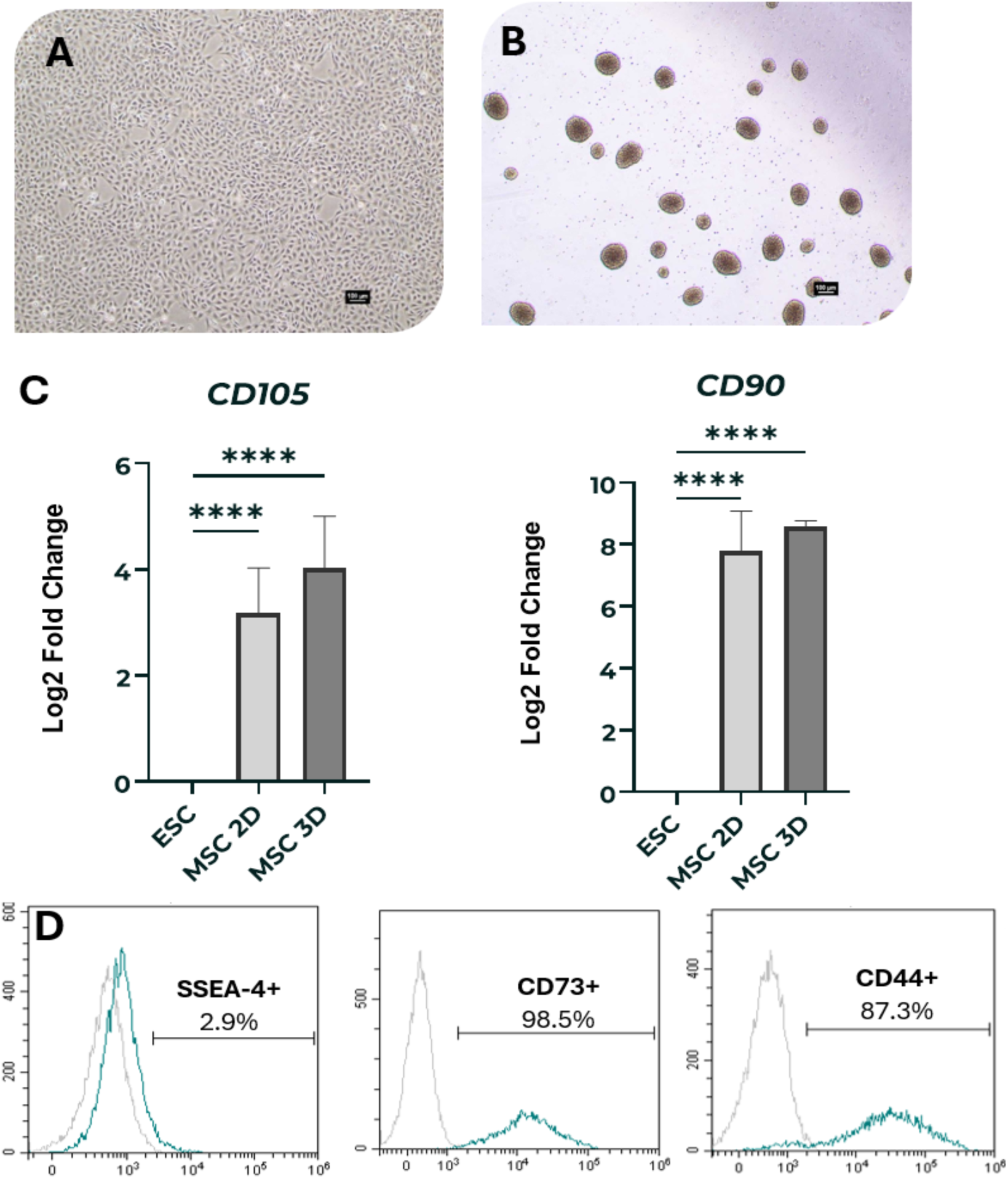
Phenotypic confirmation of bovine embryonic stem cell–derived mesenchymal stem cells in two-and three-dimensional culture. Bovine induced mesenchymal stem cells (iMSCs) were evaluated following expansion in both two-dimensional adherent culture and three-dimensional culture formats. Representative cell morphology is shown for iMSCs maintained under 2D (A) and 3D conditions (B). Molecular characterisation by RT–qPCR assessed expression of mesenchymal stem cell–associated markers relative to undifferentiated embryonic stem cells (n=3, ****=p<0.0001). Flow cytometric analysis was used to assess surface marker expression and confirm loss of pluripotency-associated antigens following differentiation, alongside acquisition of canonical mesenchymal surface markers (D) documenting the phenotypic characteristics of iMSCs expanded under planar and three-dimensional culture conditions. Scale bar = 100 µm.

**Figure 5.**
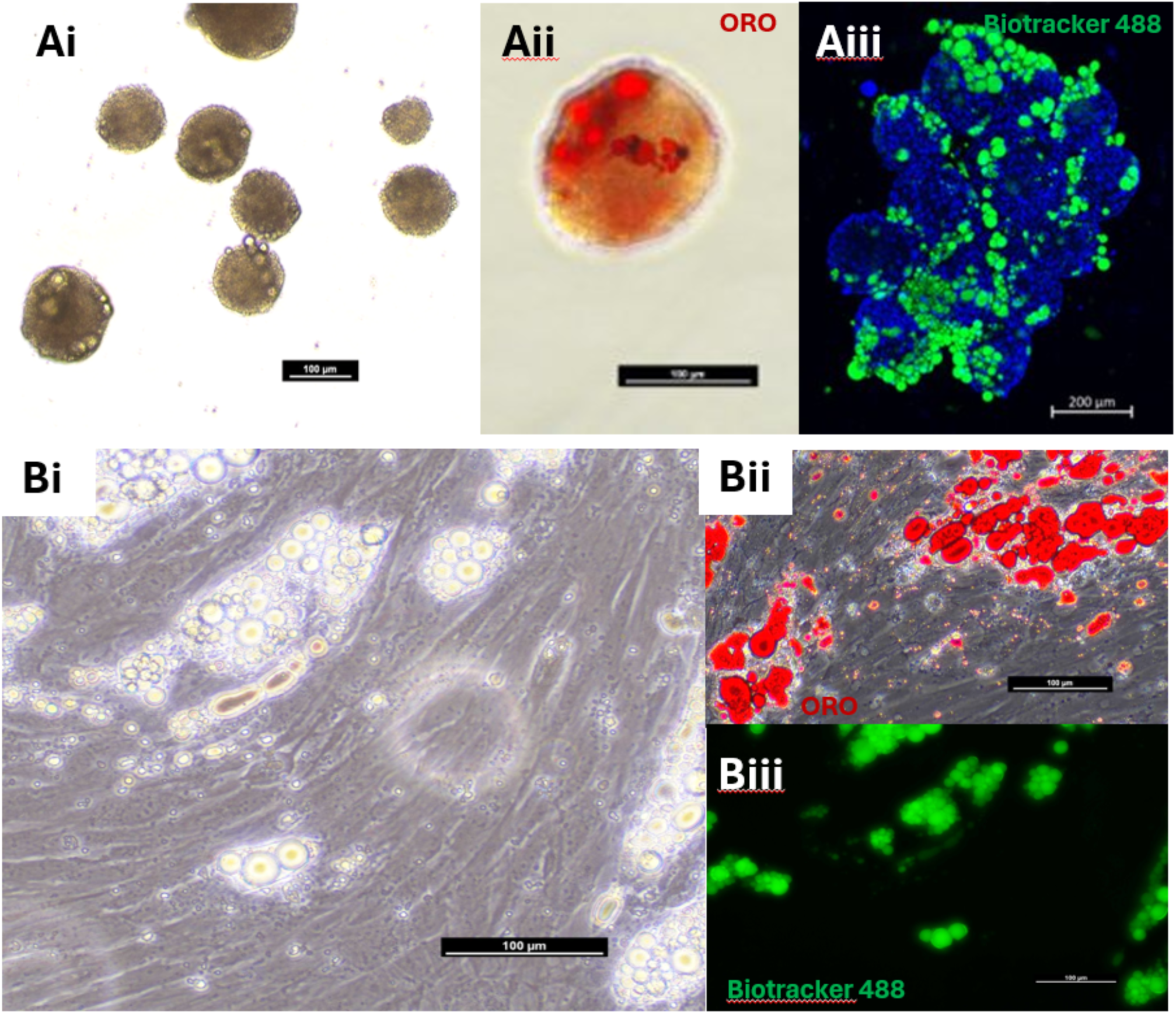
Adipogenic differentiation of bovine embryonic stem cell–derived mesenchymal stem cells in two and threedimensional culture. Bovine induced mesenchymal stem cells (iMSCs) were subjected to adipogenic differentiation under suspension and adherent culture conditions. Representative morphology of differentiated iMSCs is shown following 14 days of induction in suspension culture and 6 days of induction in adherent culture (A,B). Intracellular lipid accumulation was visualised by brightfield microscopy and confirmed using Oil Red O staining and fluorescent lipid labelling with Biotracker dye (i–iii), demonstrating adipogenic differentiation in both culture formats. Scale bar = 100 µm.

**Figure 6.**
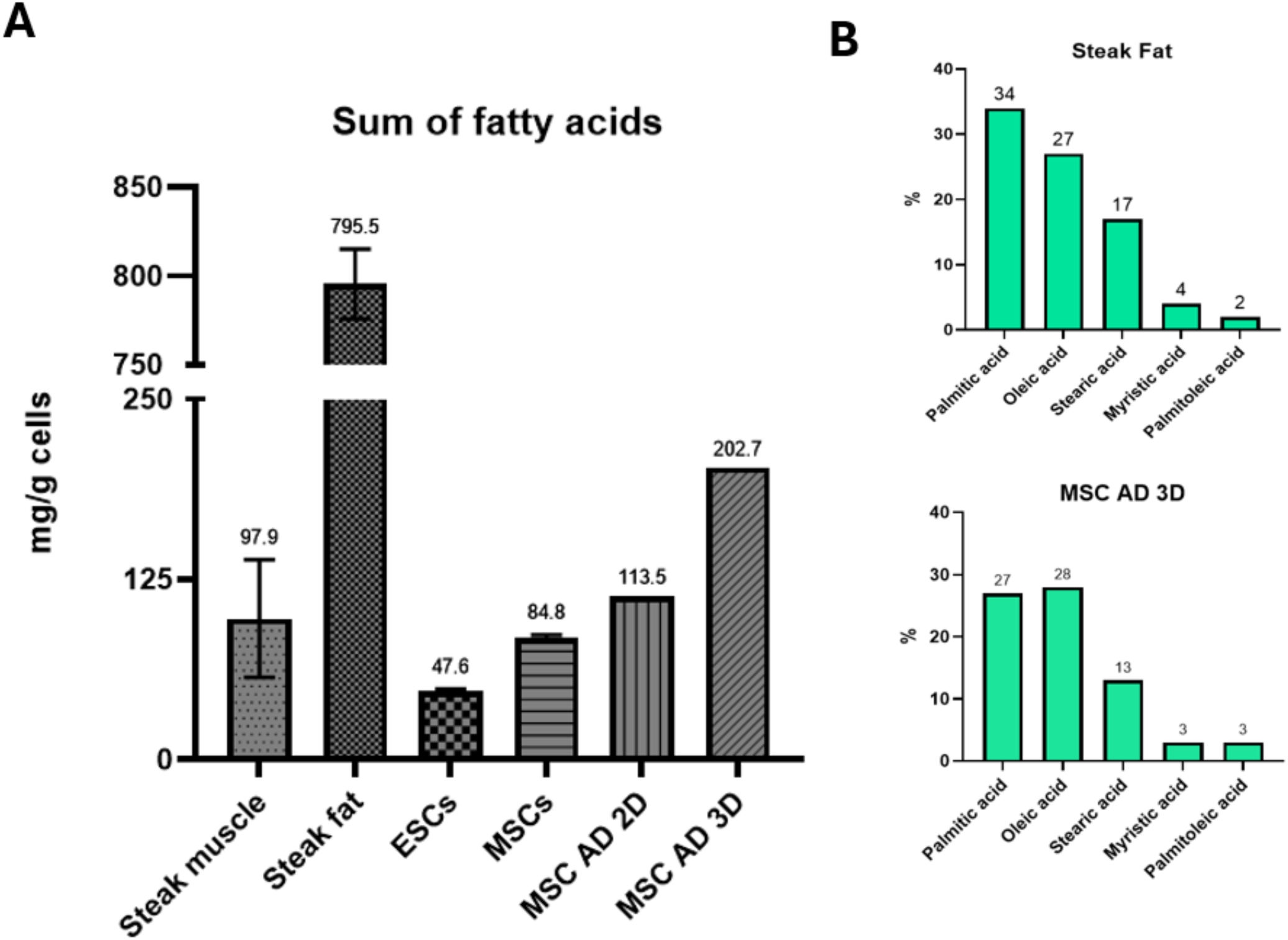
Fatty acid abundance and composition in differentiated bovine mesenchymal stem cell cultures. Fatty acid profiles were analysed in differentiated and undifferentiated bovine cell populations and compared with conventional bovine tissue samples. Total fatty acid abundance is shown for differentiated adipocyte cultures relative to undifferentiated embryonic stem cells and mesenchymal stem cells, alongside supermarket beef muscle and fat samples (A). Fatty acid composition is presented as the relative abundance of the five most prevalent fatty acids, enabling comparison between conventional steak fat and 3D differentiated iMSC adipocyte cultures (B).

**Figure 7.**
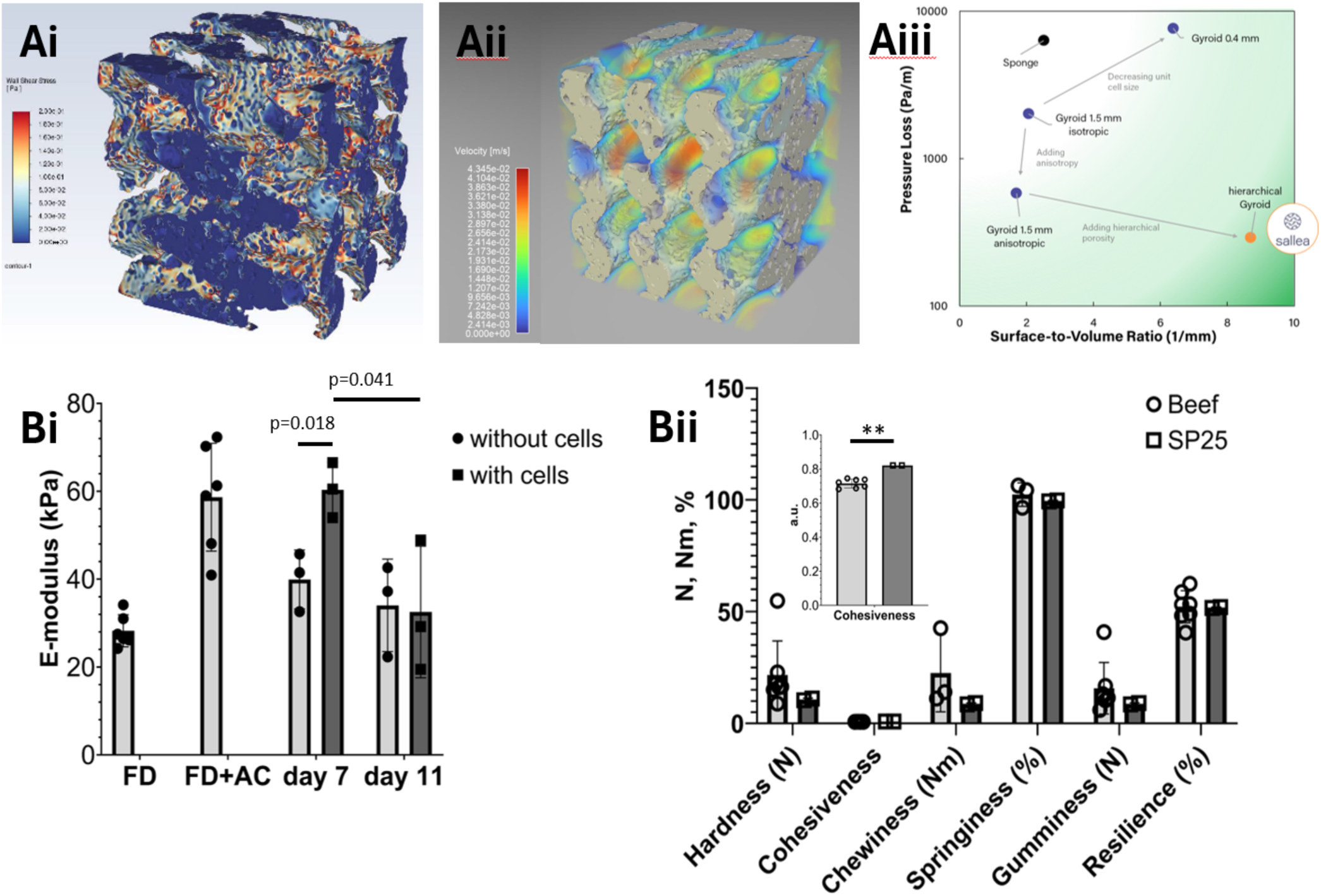
Shear environment and mechanical properties of edible protein-based scaffolds used for cell culture. A) Shear stress characterization of sallea scaffolds. i) Wall shear stress, ii) Flow velocity through the channels, iii) Pressure drop loss as a function of scaffold architecture. B) Mechanical characterization of sallea scaffolds. i) E-modulus quantification of freeze dried (FD), autoclaved (FD+AC) and cultured scaffolds with and without cells after 7 and 11 days, ii) TPA comparison between beef and soy-based scaffolds (SP25). No statistical differences were observed except for cohesiveness (p=0.0009) although scaffolds showed lower average values for all parameters (n=7 for beef and n=2 for SP25).

**Figure 8.**
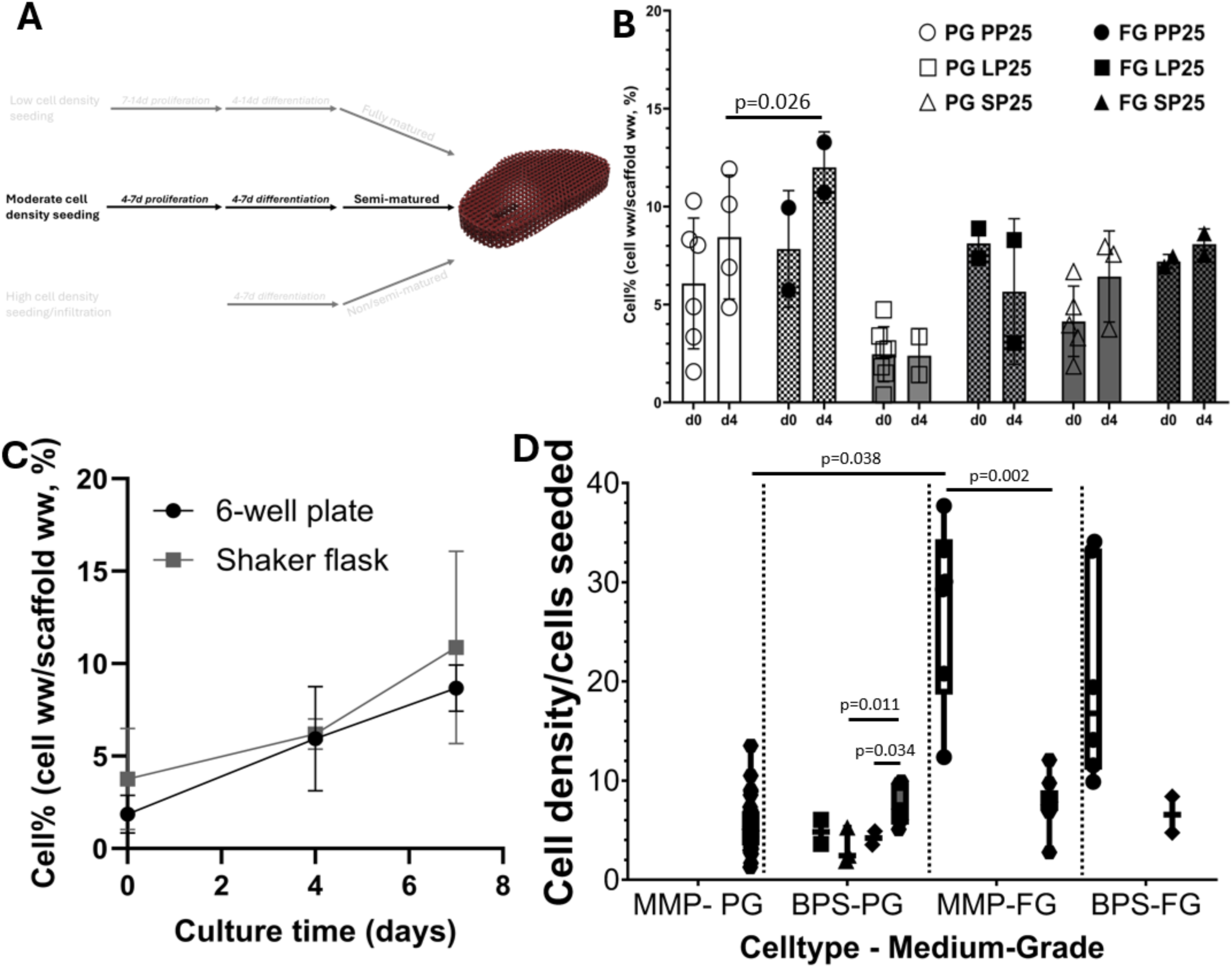
Evaluation of scaffold-based culture conditions across medium composition, scaffold material, and culture format. (A) Bioprocess development for scaffold cultures focusing on moderate cell density seeding (graphic produced on Biorender.com) (B) Scaffolds were compatible with both Pharma-Grade (PG) and Food-Grade (FG) medium and pea-based (PP25), lentil-based (LP25) or soy-based (SP25) scaffolds with cells showing no difference in growth response in either medium composition. C) Scaling prototyping showed that comparable growth can be obtained in 125mL shaker flasks and 6 well cultures on a rocker (30rpm). D) ML analysis of a partial data set revealed optimization parameters for bioprocess development (●1M cells seeded, ▪ 1.7M cells seeded, ▴ 2M cells seeded, ◆ 3.5M cells seeded,⬣5M cells seeded. MMP-PG = murine muscle progenitors in Pharma-Grade medium, BPS-PG = bovine primary satellites in Pharma-Grade medium, MMP-FG = murine muscle progenitors in Food-Grade medium, BPS-FG = bovine primary satellites in Food-Grade medium.

Cell attachment efficiency (ηₐ) was calculated as the ratio of the number of cells measured after 1 h of attachment (Cₓₙ) to the initial number of cells seeded (Cₓ₀), providing a measure of the proportion of cells successfully adhering to the scaffold (ηₐ = Cₓₙ / Cₓ₀). The fold increase (Fᵢ) in cell number was determined by dividing the cell concentration at a given time point (Cₓₙ) by the initial cell concentration at day 0 (Cₓ₀), reflecting overall cell expansion during culture (Fᵢ = Cₓₙ / Cₓ₀). The population doubling number (PD) was calculated as log(Cₓₙ / Cₓ₀) divided by log(2), in accordance with De La Garza-Rodea et al. (2012).

Cell coverage of scaffolds was defined as the percentage of scaffold mass occupied by cells and was calculated as the ratio of cell wet weight to scaffold wet weight. Specifically, cell wet weight (as determined from the standard curve and corrected for the proportion of scaffold analysed) was divided by the total scaffold wet weight to yield the percentage cell coverage.

### iMSC culture

Bovine ESC-derived mesenchymal stem cells (iMSCs; RT Bovine iMSC_5A, Holstein/Female; RT Bovine iMSC-8A, Angus x Simmental/Male) were maintained in adherent culture on tissue culturetreated plates or flasks (StarLab, CC7682-7506; Avantor, 734-2313) coated with 0.2% (w/v) gelatin (SigmaAldrich, G1393). Cells were cultured in a proprietary, serumfree MSC proliferation medium (Roslin Technologies, UK), with medium exchange every 2–3 days. The exact media formulation is proprietary and supplied by the manufacturer.

For suspension culture, bovine iMSCs were maintained in MSC proliferation medium in an Ambr® 250 unbaffled stirred-tank vessel (Sartorius, 001-5A33). Ambr® 250 run parameters were maintained at 38.5 °C with a deadband of 0.5, pH was maintained at 7.4 with a deadband of 0.05, dissolved oxygen was maintained at a minimum of 30% by controlled sparging of air, equating to a minimum of 6.3% pure oxygen, stirring speed was maintained in the range from 100 – 110 rpm, the initial working volume was 100 mL and cultures were maintained by fed-batch process up to a volume of 200 mL. Suspension culture methods were additionally validated in ultralow attachment 6well plates (Corning, 3471) and 125 mL shaker flasks (Corning, 431143) cultured on an orbital shaker platform (orbital diameter 19 mm) (Thermo Fisher Scientific, 88881102) set to 90–110 rpm, using the same proliferation medium. Cell seeding density was maintained at 5 × 10⁵ cells mL⁻¹ across all suspension culture systems.

Cells were passaged every 4–7 days when confluency of approximately 80% was reached and dissociated to single cells using TrypLE™ Select Enzyme (Gibco, 12563029). For adherent culture, cells were reseeded at 1 × 10⁴ cells cm⁻², while suspension cultures were reseeded at 5 × 10⁵ cells mL⁻¹. Cells used for experiments were between passages 3 and 5.

For adipogenic differentiation, bovine iMSCs were expanded in adherent or scaffoldbased culture until reaching confluency (approximately 90-100%). Cells were then cultured in a proprietary, serumfree adipogenic induction medium (Roslin Technologies) for 4 days, followed by a proprietary adipogenic maintenance medium (Roslin Technologies). Adipogenic maintenance was continued for an additional 3 days for adherent and scaffold cultures, or 10 days for suspension cultures. The media formulations are proprietary and supplied by the manufacturer.

All iMSC cultures were maintained in a humidified incubator or bioreactor at 38.5 °C with 5% CO₂. Routine sterility testing was performed throughout the study, and all cultures were confirmed to be free of mycoplasma contamination using a VenorGEM qEP Mycoplasma Detection Kit assay (Cambio, 11-9250).

### Bovine satellite cell and murine cell culture

Primary bovine satellite cells (Opo-Moo-M18; Opo Bio, New Zealand) were cultured according to the manufacturer’s protocol. Briefly, cells were seeded at 3 × 10^3^ cells cm⁻² in T300 tissue culture flasks (TPP, 90301) in proliferation medium containing 20% fetal bovine serum (Sigma-Aldrich, F7524) and 5 ng mL⁻¹ basic fibroblast growth factor (bFGF; Sigma-Aldrich, GF446). Cells were maintained until reaching 70–80% confluency before harvesting by enzymatic dissociation using TrypLE™ Select Enzyme (Gibco, 12563029).

Murine muscle precursor cells (C2C12, ATCC) were seeded at 1,500 cells cm⁻² in T300 tissue culture flasks (TPP, 90301) and cultured in proliferation medium containing 10% fetal bovine serum (Sigma-Aldrich, F7524). Cells were harvested at 80–90% confluency using TrypLE™ Select Enzyme.

Primary bovine and murine cell cultures were maintained at 37 °C with 5% CO₂ in a humidified incubator.

### Scaffold generation

Porous scaffolds were fabricated using a negative salt templating approach. Salt templates were produced using stereolithography (Prusa SL1S) with proprietary ink based on NaCl particles. Following printing, the salt templates were washed and underwent a proprietary thermal treatment to remove the binder. These templates were then used as sacrificial molds for scaffold generation.

High-protein dispersions were prepared by mixing 25% (w/v) plant protein powders (pea (Bulk, 000273994), lentil (HSN, 220201C3), or soy (myvegan, P410615528); in a proprietary liquid medium (sallea AG, Switzerland). Protein dispersions were prepared in a food-safe environment and mechanically infiltrated into the printed salt templates using a 10mL syringe.

Following infiltration, the protein dispersions were cured within the salt templates for 60 min. The sacrificial salt templates were subsequently removed by dissolution in tap water at room temperature, yielding free-standing porous protein scaffolds. Scaffolds were then freeze-dried for up to 24h with a minimum of 8h using Home Freeze Dryer (Harvest Right). Initial freeze was performed at-17°C and the final drying was performed at 51°C. Scaffolds were stored dry at room temperature until use.

The resulting scaffolds were cylindrical in geometry, with an approximate height of 10 mm and diameter of 7 mm.

### Pre-culture scaffold preparation and cell seeding

Prior to cell seeding, dried scaffolds were placed in a heat-resistant glass beaker and covered with aluminum foil before sterilization by autoclaving. Scaffolds were sterilized using a custom solids dry autoclave cycle at 121 °C for 15 min, followed by a 16-min drying phase. After sterilization, scaffolds were allowed to cool under sterile conditions before further handling.

Autoclaved scaffolds were transferred with sterile tweezers to 24-well plates (TPP, 92124) pre-conditioned by incubation with 100 µL of undiluted FBS for 1 h at 37 °C (C2C12 and Opo-Moo-M18) or a 1% (w/v) bovine serum albumin (BSA) solution (Sigma-Aldrich, A1470) for 1 h at 38.5 °C (iMSCs). Excess solution was removed immediately prior to cell seeding.

For seeding, bovine iMSCs were prepared as a concentrated single-cell suspension at a density of 5 × 10⁶ cells per 80 µL. C2C12 and Opo-Moo-M18 cells were prepared at a concentrated single-cell suspension at a density between 1 and 5 × 10⁶ cells per 80 µL. A volume of 80 µL cell suspension was carefully applied directly onto each scaffold, corresponding to 5 × 10⁶ cells per scaffold. Seeded scaffolds were incubated statically for 1 h at 37 °C (C2C12 and Opo-Moo-M18) or 38.5 °C (iMSCs) to allow initial cell attachment.

Cells were seeded onto scaffolds either immediately following thawing of cryopreserved cell stocks (Figure 9) or following expansion in adherent culture prior to seeding (Figure 10). Subsequent scaffold culture was performed as described in the relevant experimental sections.

**Figure 9.**
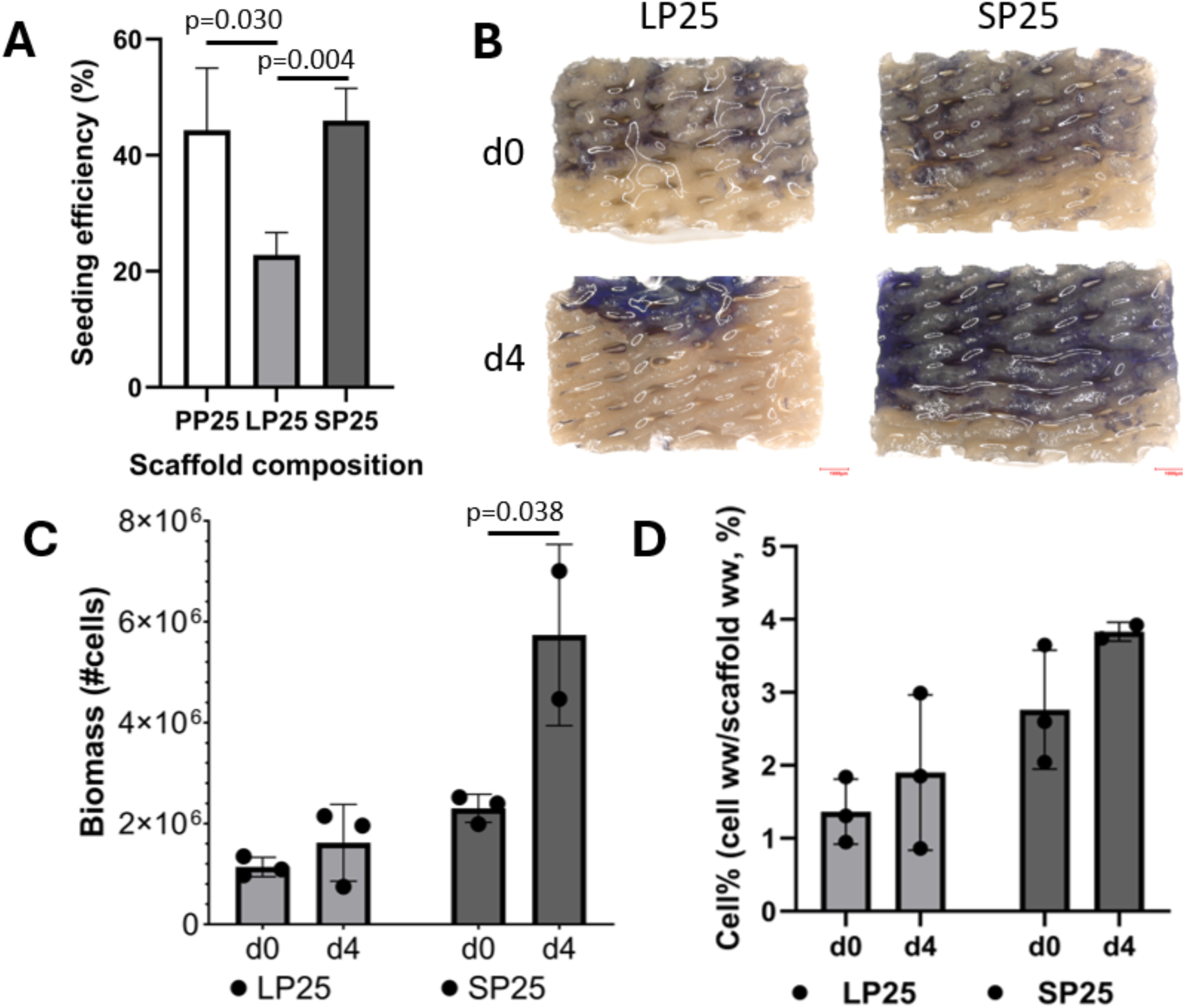
Compatibility of bovine induced mesenchymal stem cells with plant-based scaffold materials. Bovine induced mesenchymal stem cells (iMSCs) were evaluated for attachment and short-term growth on plant-derived scaffolds composed of pea-(PP25), lentil-(LP25), and soy-based protein (SP25) formulations. Initial seeding efficiency of iMSCs on the different scaffold materials is shown to compare early cell attachment across scaffold types (A). Cell distribution on lentil-and soy-based scaffolds following culture was visualised by MTT staining (B). Quantitative analysis of biomass accumulation and cell coverage was performed using MTT-based measurements to describe differences in total biomass and spatial distribution between scaffold materials (C,D)

**Figure 10.**
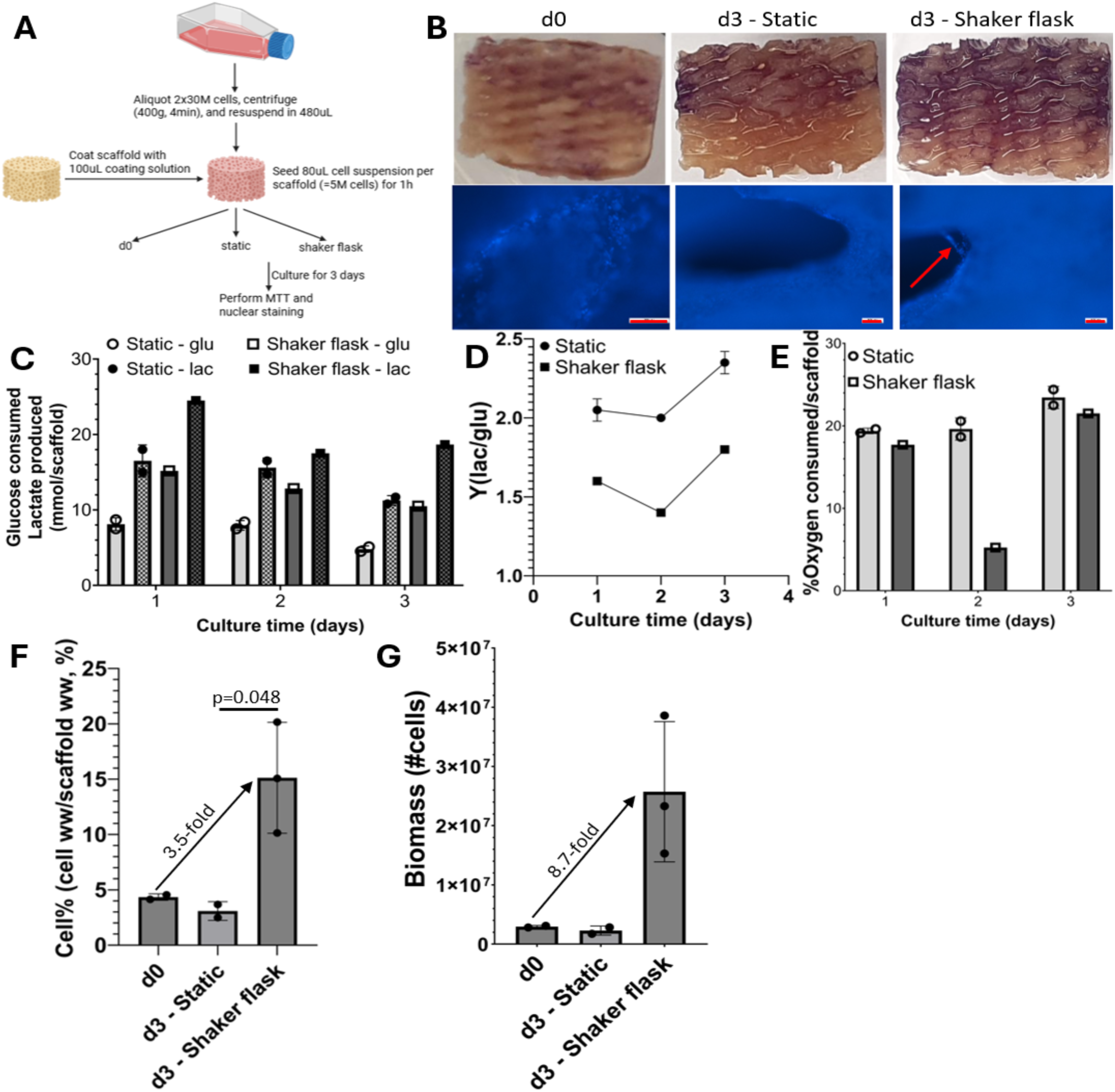
Dynamic and static scaffold-based culture of bovine induced mesenchymal stem cells. Bovine induced mesenchymal stem cells (iMSCs) were cultured on soy-based scaffolds under static and dynamic conditions to assess cell distribution, metabolic activity, and biomass accumulation. A schematic of the scaffold culture configuration and experimental strategy is shown (A). Following short-term culture, macroscopic and microscopic cell distribution on scaffolds was visualized by MTT staining and nuclear counterstaining, with representative images showing cell localisation and pore-spanning structures within the scaffold architecture (B). Metabolic activity during culture was assessed by measuring glucose consumption, lactate production, and oxygen uptake on a per-scaffold basis (C–E). Quantitative MTT-based analysis was used to estimate scaffold-associated cell coverage and total cell number following dynamic culture (F,G). Scale bar = 100 µm.

All scaffold handling and cell seeding steps were carried out using aseptic technique within a class 2 biological safety cabinet.

### Scaffold culture

#### Screening

Following cell seeding, soy-based (SP25) or lentil-based (LP25) scaffolds were transferred to ultra-low attachment 6-well plates (Corning, 3471), with one scaffold per well, and cultured in 3 mL of proprietary, serum-free MSC proliferation medium (RBDM1; Roslin Technologies, Edinburgh, UK). Plates were placed on a Benchmark Orbi-Shaker BT3001 rocking platform operating at 30 rpm and maintained for 4 days in a humidified incubator at 38.5 °C with 5% CO₂. Culture medium was replaced daily.

#### Site validation

Following cell seeding, soybased scaffolds (SP25) were cultured either in ultralow attachment 6well plates (Corning, 3471) or 125 mL shaker flasks (Corning, 431143), with three scaffolds per flask. Scaffolds cultured in 6well plates were maintained in 3 mL of MSC proliferation medium (Roslin Technologies) per well, with daily complete medium changes.

For shaker flask cultures, scaffolds were maintained in 35 mL of MSC proliferation medium on an orbital shaker (orbital diameter 19 mm) (Thermo Fisher Scientific, 88881102) operating at 70 rpm. On day 2 of culture, 50% of the culture medium was exchanged for an equal volume of fresh medium. All cultures were maintained for a total of 3 days in a humidified incubator at 38.5 °C with 5% CO₂.

### Scaffold processing and viability analysis

At the end of the culture period, scaffolds were removed from culture and bisected longitudinally using a sterile scalpel. One half of each scaffold was used for metabolic activity assessment by MTT staining and quantification, while the remaining half was reserved for nuclear staining and imaging as described below. Scaffold halves were briefly placed on absorbent paper to remove surface liquid before weighing to determine wet weight (ww) using an analytical balance (Veritas, LW203I).

### MTT assay and biomass quantification

Scaffold halves allocated for metabolic analysis were incubated in 0.5 mL of 0.5 mg mL⁻¹ MTT solution (Sigma Aldrich, M5655) prepared in phosphate-buffered saline. Incubations were performed for 40 or 60 min at 38.5 °C under static conditions. Following incubation, scaffolds were imaged using a Keyence VHX-S660E stereomicroscope to document spatial distribution of formazan formation.

To solubilize the reduced MTT product, scaffold halves were submerged in 0.5 mL acidic ethanol 0.01 M HCl (Fisher Scientific, 12943544) in absolute ethanol (Fisher Scientific, 16685992) and incubated overnight at 4 °C. The resulting eluates were transferred to a transparent 96well plate, and absorbance was measured at 600 nm using a plate reader (PerkinElmer, VICTOR® Nivo™).

Total biomass and the ratio of cell wet weight to scaffold wet weight (cell%, expressed as ww/ww) were calculated using standard curves generated from defined bovine iMSC numbers. Cell pellets of known cell numbers were prepared, centrifuged at 400 x *g* to remove excess liquid and weighed to determine pellet wet weight. Pellets were incubated under identical conditions and for the same duration as scaffold samples during the MTT assay and dye solubilization steps. Standard curves were generated for the following relationships: number of cells versus pellet wet weight, number of cells versus MTT absorbance, and pellet wet weight versus MTT absorbance.

### Metabolic analysis

Culture supernatant samples were collected at the end of the indicated culture periods and clarified by brief centrifugation 400 × *g* for 2 min) prior to analysis. Supernatant pH, dissolved oxygen, dissolved carbon dioxide, and metabolite concentrations were analyzed using a MetaFLEX analyzer (Beckman Coulter/ViCell, Brea, CA, USA).

Metabolite analysis included quantification of glucose and lactate according to the manufacturer’s protocols. The analyzer was calibrated prior to sample measurement using manufacturersupplied standards and quality controls.

### RT-qPCR

Cell pellets were collected by centrifuging samples for 4 min at 400 × *g*, washing with DPBS (Gibco, 14190) and snap freezing at-80⁰C. RNA was extracted using the RNeasy Mini kit (Qiagen, 74136), quantified by Nanodrop, and reverse transcribed using AffinityScript cDNA synthesis kit (Agilent, 200436) following manufacturer’s instructions. qPCR reactions were prepared with 1x PowerSYBR green Mastermix (Applied Biosystems, 4367659), custom primers for the relevant target (IDT) at optimized concentrations of 50-300nM, and 10ng sample cDNA per reaction and analyzed in a qPCR machine (Applied Biosystems, Quantstudio3). Sample results were normalized to a housekeeping gene HMBS and to a calibrator sample according to the delta Ct method (Schmittgen & Livak, 2008). Relative gene expression data was compared between test samples and undifferentiated controls as log2 fold change and analyzed by one-way ANOVA (GraphPad Prism).

### RNA-sequencing

Total RNA was extracted from samples and quantified as described above. Library preparation and sequencing were performed by an external service provider (GENEWIZ, Azenta Life Sciences). RNAsequencing libraries were prepared from RNA isolated from ESCs, MSCs, and MP samples using the TruSeq Stranded mRNA Library Preparation Kit (Illumina). Libraries were sequenced on an Illumina NovaSeq 6000 platform to generate 150 bp pairedend reads, with an average depth of approximately 30 million reads per sample.

Raw sequencing reads were quality-checked using FastQC (v0.12.1). Adapter sequences and low-quality reads were removed using Cutadapt (v3.2) (Martin, 2011). Filtering criteria included removal of reads containing adapter sequences, reads with a Phred quality score <30, reads shorter than 50 bp after trimming, and reads with undetermined base calls (N) at the 5′ or 3′ ends.

Filtered reads were aligned to the *Bos taurus* reference genome (NCBI ARS-UCD2.0; assembly GCF_002263795.3) using STAR (v2.7.11a) (Dobin et al., 2013) with default parameters. Gene-level read counts were generated using featureCounts (v2.0.6) (Liao et al., 2014).

Differential gene expression analysis was performed using edgeR (v4.2.2) (Chen et al., 2025). Genes with an adjusted *P*-value < 0.05 and an absolute log₂ fold change ≥ 2 were considered significantly differentially expressed.

### Flow cytometry

Singlecell suspensions were prepared as described above and incubated in cell staining buffer consisting of Dulbecco’s phosphatebuffered saline (DPBS; Gibco, 14190-144) supplemented with 2% (v/v) chicken serum (Gibco, 16110). Cells were stained with a fixable viability dye (Invitrogen, L34963) and fluorophoreconjugated primary antibodies against CD44 (BioLegend), CD73 (Life Technologies), and SSEA4 (BioLegend). Antibody incubations were performed for 40 min at ambient room temperature (20-23 °C) protected from light.

Following primary antibody staining, cells were washed with staining buffer and incubated with an AlexaFluor™ 647conjugated donkey antiIgG secondary antibody (Invitrogen, A21448) at a dilution of 1:1000 for 30 min at ambient room temperature (20-23 °C) protected from light. Cells were then washed again with staining buffer and resuspended in 100 µL for analysis.

Flow cytometric analysis was performed using a CytoFLEX™ 5 flow cytometer (Beckman Coulter). Compensation was carried out using fluorescence minus one (FMO) and unstained controls, and data were acquired and analyzed using CytExpert™ software (Beckman Coulter). Gating strategies were based on forward and side scatter to exclude debris, followed by viability gating and markerspecific fluorescence thresholds.

### Staining and imaging

Adipogenic differentiation was assessed by lipid staining. For Oil Red O (ORO) staining, cultures were fixed with 4% (w/v) paraformaldehyde (Thermo Fisher Scientific, J61889) and rinsed with 60% isopropanol (SigmaAldrich, I9516). Fixed samples were then incubated with a 60% ORO solution (SigmaAldrich, O0625) for 10 min at room temperature, followed by washing with sterile distilled water to remove excess stain prior to imaging.

Alternatively, lipid accumulation in live cells was visualized by incubation with a fluorescent lipid dye. Live cultures were incubated with BioTracker™ 488 lipid dye (Merck, 3CT144) for 30 minutes at 38.5 °C according to the manufacturer’s instructions before imaging.

For nuclear staining, live cell samples were incubated with NucBlue™ nuclear stain (Invitrogen, R37605) for 30 min at 38.5 °C prior to imaging.

Imaging was performed using an inverted fluorescence microscope equipped for brightfield and fluorescence imaging with 488 nm and 360 nm filter sets (Nikon Eclipse TS2). Where indicated, higher-resolution imaging was carried out using confocal laser scanning microscopy (Zeiss LSM 880). For all imaging modalities, acquisition and exposure settings were kept constant between experimental samples and corresponding controls.

### Fatty acid analysis

Cell samples were collected by centrifugation at 400 × *g* for 4 min, washed once with Dulbecco’s phosphatebuffered saline (DPBS; Gibco, 14190144), and snapfrozen at −80 °C. Control tissue samples representing conventional beef fat and muscle were obtained from a commercially available beef sirloin steak (Sainsbury’s, UK), portioned into sterile tubes, and snapfrozen at −80 °C.

Frozen samples were lyophilized using a benchtop freeze dryer (Frozen in Time, Lablyo Mini) prior to weighing. Lyophilized samples were subsequently analyzed for fatty acid composition and quantification by gas chromatography at an external analytical service provider (Vitas Analytical Services, Oslo, Norway).

## Results

### Generation and characterization of bovine iMSCs in 2D and 3D culture

To establish a scalable bovine MSC source with adipogenic potential, pluripotent RT bovine embryonic stem cells (ESCs) were differentiated through an intermediate mesodermal progenitor (MP) stage to an induced mesenchymal stem cell (iMSC) state (Figure 2). This staged differentiation strategy generated a cryo-preservable mesodermal progenitor intermediate population and a downstream iMSC population suitable for expansion and functional evaluation.

To assess transcriptional changes throughout the differentiation pathway, RNAsequencing was performed on ESCs, MPs, and iMSCs. Unsupervised clustering revealed that each differentiation stage formed a transcriptionally distinct population, indicating clear transitions in gene expression programs during lineage progression (Figure 3). ESCs derived from three individual animals clustered tightly by cell type, while MP and iMSC populations derived from two ESC lines (BEIH5 and BEIH8) formed separate, stagespecific clusters with high similarity between biological replicates (Figure 3, Supplementary Figure 1). These data demonstrate that differentiation from ESCs to iMSCs is associated with reproducible and welldefined transcriptional remodeling rather than gradual or heterogeneous drift.

Following differentiation, bovine iMSCs were evaluated for their capacity to expand under both twodimensional (2D) and threedimensional (3D) culture conditions. In adherent 2D culture, iMSCs formed a monolayer and exhibited an elongated spindleshaped morphology characteristic of mesenchymal stem cells (Figure 4A). In parallel, iMSCs were successfully expanded in 3D suspension culture within stirredtank bioreactors, where they selfassembled into multicellular aggregates with diameters below 400 µm (Figure 4B). These observations indicate that bovine iMSCs can be propagated in both planar and suspension formats without loss of viability or gross morphological abnormalities. To confirm acquisition of a mesenchymal phenotype, expression of canonical MSC markers was assessed following expansion in both culture formats. After seven population doublings in suspension culture and ten population doublings in adherent culture, iMSC populations showed significantly increased expression of the MSC markers CD90 and CD105 compared to undifferentiated ESCs, as measured by RTqPCR (p < 0.0001, n = 3) (Figure 4C). These results confirm successful transition from a pluripotent to mesenchymal transcriptional state and demonstrate that marker expression is maintained during early expansion.

Cell type purity within expanded iMSC banks was further assessed by flow cytometry. After 12 population doublings in adherent culture, more than 85% of cells exhibited coexpression of the MSCassociated surface markers CD73 and CD44 (Figure 4D), indicating a highly enriched mesenchymal population. Due to the lack of speciescompatible antibodies, hematopoietic exclusion markers (CD45, CD34, CD14, and HLADR) could not be evaluated; however, residual expression of the ESCassociated marker SSEA4 was detected in fewer than 3% of cells (Figure 4D), suggesting minimal persistence of undifferentiated cells. Consistent with their mesenchymal identity, iMSCs showed strong adherence to tissueculture plastic and could be readily expanded under standard adherent conditions (data not shown).

Collectively, these results demonstrate that bovine iMSCs can be reproducibly derived from pluripotent ESCs, exhibit stable mesenchymal marker expression following expansion, and are compatible with both adherent and suspension culture formats suitable for downstream bioprocessing applications.

### Adipogenesis of bovine iMSCs is possible in 2D and 3D culture

To assess whether bovine iMSCs retained functional lineage competence following expansion, cells cultured in either three-dimensional (3D) suspension culture (7 population doublings) or two-dimensional (2D) adherent culture (12 population doublings) were induced toward adipogenic differentiation. This enabled evaluation of adipogenic potential following expansion under both planar and suspension formats relevant to bioprocess development.

Following adipogenic induction, cultures in both expansion formats accumulated visible intracellular lipid droplets (Figure 5Ai, Bi). Lipid deposition was evident by brightfield imaging and was further confirmed by positive Oil Red O staining and fluorescent lipid labeling (Figure 5Aii, Bii). In contrast, undifferentiated control cultures showed no detectable lipid accumulation (Supplementary Figure 2), indicating that lipid formation was dependent on adipogenic induction rather than prolonged culture.

At the molecular level, differentiated adipogenic cultures exhibited significantly increased expression of the adipogenic late-stage marker genes *FABP4* and *PPARγ* relative to undifferentiated controls (p < 0.005, n = 3) (Supplementary Figure 3), consistent with activation of an adipogenic gene regulatory program.

To quantify lipid accumulation, fatty acid (FA) composition was measured in differentiated cultures and compared with undifferentiated cell populations. Adipogenic differentiation under 3D conditions resulted in a fourfold increase in total FA abundance relative to undifferentiated ESCs and a 2.4fold increase compared to undifferentiated iMSCs (Figure 6A), consistent with the establishment of a lipidrich phenotype.

In addition to increased lipid abundance, differentiated adipogenic cultures displayed a fatty acid composition broadly comparable to that of conventional bovine adipose tissue. Palmitic acid, oleic acid, and stearic acid were the most abundant fatty acids detected in both differentiated cultures and supermarket beef steak fat samples (Figure 6B). This similarity indicates that iMSCderived adipogenic cultures generate lipid profiles aligned with native bovine fat.

Together, these results demonstrate that bovine iMSCs retain robust adipogenic differentiation capacity following both 2D and 3D expansion, resulting in increased lipid accumulation and fatty acid compositions characteristic of bovine adipose tissue, despite current limitations in speciesspecific reagent availability for some adipogenic markers.

### Scaffold design for optimal cell culture conditions

To evaluate scaffold architectures suitable for scalable cell culture, structured protein-based scaffolds were fabricated with a highly ordered gyroid geometry and compared with conventional sponge-like scaffolds. Key physical properties relevant to cell culture performance (including pressure loss, internal surface area, and mechanical stability) were assessed.

Computational and experimental characterization showed that gyroid structured scaffolds had up to 2 x 10^-1^ Pa and exhibited a 22-fold reduction in pressure loss compared with sponge-like constructs (Figure 7Ai). This resulted in a uniform flow pattern in the interior of the scaffold with a flow rate lower than 4.3 x 10^-2^ m/s, in a range suitable for cell proliferation (Figure 7Aii). In parallel, the internal surface area of the gyroid scaffolds increased approximately 3.5-fold relative to sponge controls (Figure 7A). These combined changes indicate substantially improved internal mass transfer characteristics while increasing available surface area for cell attachment.

Mechanical performance of the scaffolds was evaluated during extended exposure to culture conditions. Scaffolds retained structural integrity for up to 11 days in culture medium at 37°C, both in the presence and absence of cells, indicating that the material and architecture were stable under biologically relevant conditions.

Quantitative mechanical testing further showed that the elastic modulus (E-modulus) of soy-based scaffolds fell within a range of 35-60 kPA, previously reported to support growth and differentiation of mesoderm-derived cells (Figure 7Bi) (Pavan et al., 2020; Ward et al., 2009). These measurements indicate that scaffold stiffness is compatible with MSC culture without imposing excessive mechanical constraint.

To benchmark scaffold textural properties against conventional meat tissues, texture profile analysis was performed. No statistically significant differences were observed between hydrated soybased scaffolds without cells and beef samples across most measured parameters, with the exception of increased cohesiveness in the scaffold material (Figure 7Bii). This result indicates partial overlap between scaffold mechanical properties and those of native muscle tissue, while highlighting a distinct difference in cohesiveness.

Together, these results demonstrate that gyroid-structured protein scaffolds provide favorable mass-transfer properties, mechanical stability, and tissue-relevant textural characteristics, supporting their suitability for subsequent high-density cell culture applications.

### Experimental conditions for seeding research cell lines on plant scaffolds

To determine suitable experimental seeding and culture conditions on plantderived scaffolds, multiple culture configurations were evaluated with research cell lines, including variations in scaffold material, medium grade, seeding density, and culture format (Figure 8). Murine and primary bovine progenitor cells were initially used as a screening model to compare scaffold materials and media formulations under controlled conditions.

During a four-day culture period, murine progenitor cells were seeded onto pea-based (PP25), soy-based (SP25) and lentil-based (LP25) scaffolds under rocking conditions in low attachment 6-well plates and accumulated measurable biomass in both food-grade (FG) and pharmaceutical-grade (PG) basal media (Figure 8B). Comparative analysis of scaffold materials showed that murine progenitor cells preferentially accumulated biomass on pea-based scaffolds relative to lentil-or soy-based scaffolds (Figure 8B), indicating scaffold-specific differences in support of cell attachment and growth. Cultures maintained in FG medium achieved a mean biomass content of 12.5%, compared with 8% in PG medium (Figure 8B). These results demonstrate that medium formulation influences biomass accumulation under otherwise identical culture conditions.

Based on these outcomes, both pea and soy-based scaffold types were screened in subsequent experiments with bovine iMSCs.

To assess scalability and transferability between culture formats, scaffold cultures were compared between 6-well rocking conditions and 125 mL shaker flask cultures. Cultures maintained in dynamic rocking conditions showed increasing cell growth over 7 days (Figure 8C). No statistically significant differences in final cell numbers were observed between the two systems (Figure 8C), indicating that scaffold-based cultures could be transferred from static or semi-static formats to more dynamic, scalable culture conditions without loss of cell yield.

Biomass production efficiency was further evaluated by analyzing the ratio of final cell density to initial seeding density using a machine learning–based analytical approach (LABMaiTE GmbH). This analysis revealed that, in FG medium, both murine progenitor cells and bovine satellite cells achieved higher biomass production efficiency when seeded at lower initial densities (Figure 8D). In contrast, bovine cells cultured in PG medium showed higher efficiency at increased seeding densities. These results indicate that optimal seeding density is both cell type and medium dependent on soy-based scaffolds.

Collectively, these data demonstrate that scaffold material, medium grade, and seeding density significantly influence biomass accumulation and production efficiency, and that optimized scaffold cultures are transferable between laboratory-scale and dynamically mixed culture systems.

### Comparison of bovine iMSC growth on soy and lentil-based scaffolds

To assess scaffold compatibility for bovine iMSC culture, cell attachment and shortterm expansion were compared across plantderived scaffold materials. Initial screening indicated that bovine iMSCs attached more efficiently to pea and soybased scaffolds than to lentilbased scaffolds (Figure 9A), suggesting similar materialdependent differences in early cell–scaffold interactions (Figure 8B).

To further compare scaffold performance, bovine iMSCs were cultured on lentil and soybased scaffolds for four days under identical conditions. Cells cultured on soybased scaffolds exhibited a more homogeneous spatial distribution throughout the scaffold compared with those on lentilbased scaffolds (Figure 9B). Quantitative analysis showed significantly higher total cell number on soybased scaffolds after four days of culture (Figure 9C), together with an increased ratio of cell wet weight to scaffold wet weight (Figure 9D), indicating greater biomass accumulation than that on lentil-based scaffolds.

In addition to culture in low volume static and rocked 6-well plate culture formats, the short-term compatibility of bovine iMSC–scaffold constructs was evaluated in dynamic systems. iMSC-seeded scaffolds were successfully maintained during short-term culture in perfusion and TideMotion bioreactor systems with cell attachment and without observable loss of scaffold integrity (data not shown), supporting the transferability of the soy-based scaffold–cell combination to more dynamic culture environments.

Taken together, these results demonstrate that soy-based scaffolds provide superior support for bovine iMSC attachment, distribution, and early biomass accumulation compared with lentil-based scaffolds and are compatible with both static and dynamic culture formats relevant to scalable cultivation.

### Bovine iMSC culture optimized for dynamic culture on soy-based scaffolds

To evaluate shortterm, highdensity bovine iMSC culture under conditions relevant to dynamic processing, soybased scaffolds were used in a sitevalidation experiment designed to assess reproducibility in a different laboratory environment and to compare static versus shakerflask culture over a threeday period (Figure 10A). The experiment focused on highdensity seeding and short culture durations to enable direct comparison of early attachment, distribution, metabolic performance, and biomass accumulation under dynamic mixing conditions.

After three days of culture, MTT staining indicated that iMSC coverage on soybased scaffolds under dynamic shakerflask conditions was comparable to the coverage observed previously at four days in the earlier scaffold comparison study and was increased relative to day 0 controls and statically cultured scaffolds (Figure 10). Consistent with this, nuclear staining showed regions in which cells began bridging scaffold pores under dynamic conditions, indicating progression from surface attachment toward threedimensional coverage within the scaffold architecture (Figure 10B).

Culture supernatant analysis demonstrated distinct metabolic profiles between dynamic and static conditions (Figure 10C). Shakerflask cultures showed higher glucose consumption and lactate production rates relative to static controls, alongside lower oxygen consumption per scaffold (Figure 10C, 10D, 10E). Together, these measurements are consistent with increased overall metabolic activity and greater biomass accumulation under dynamic culture conditions (Figure 10).

Quantitative MTTbased analysis further supported enhanced biomass accumulation in shaker-flask cultures. Biomass accumulation (cell wet weight/scaffold wet weight) was significantly higher under dynamic conditions than under static culture, with an average 3.5fold increase in cell wet weight fraction to approximately 15% after three days (Figure 10F). This corresponded to an approximately 8.7fold increase in biomass, with mean yields of approximately 25 × 10⁶ cells per scaffold (Figure 10F–G). These quantitative outcomes align with the higher metabolite turnover observed in dynamic cultures (Figure 10C).

Collectively, these results show that bovine iMSCs can be cultured reproducibly at high density on soy-based scaffolds under dynamic conditions, achieving rapid increases in scaffold coverage and biomass accumulation over short culture durations, with accompanying increases in measurable metabolic activity.

### Functional analysis of bovine iMSC-scaffold conditions by adipogenic differentiation

To determine whether bovine iMSCs cultured on soy-based scaffolds under dynamic conditions retained functional differentiation capacity, scaffold-cultured cells were subjected to a seven-day adipogenic differentiation protocol following high-density shaker-flask culture for 7 days (Figure 11A). This experiment was designed to assess adipogenic competence after exposure to scaffold materials and dynamic culture conditions relevant to scale-up.

**Figure 11.**
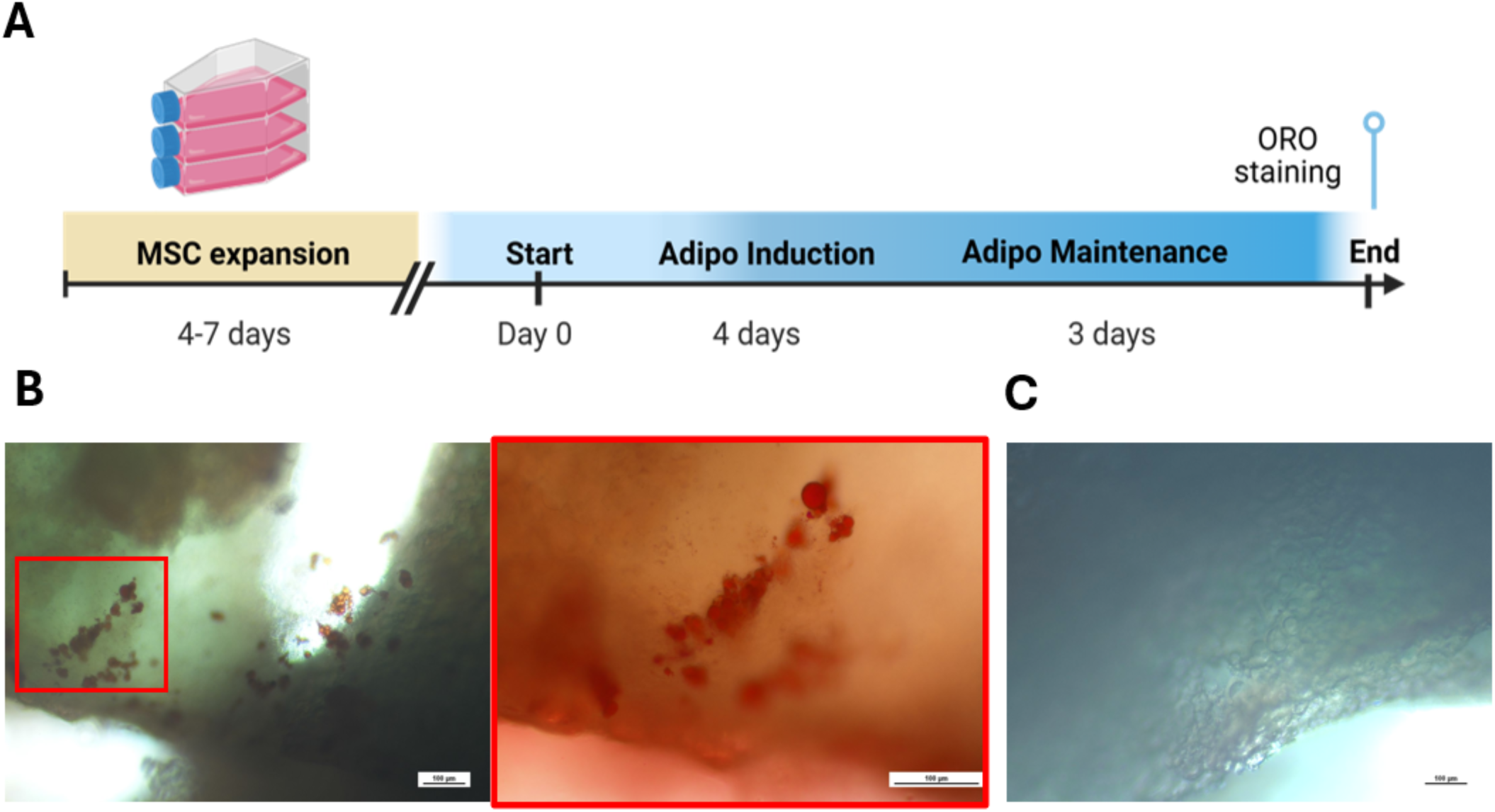
Adipogenic differentiation of bovine induced mesenchymal stem cells cultured on soy-based scaffolds. Adipogenic differentiation of bovine induced mesenchymal stem cells (iMSCs) cultured on edible protein-based scaffolds was evaluated following scaffold-based expansion. A schematic illustrating the timeline of iMSC differentiation toward adipocytes is shown (A). After culture in adipogenic induction and maintenance media, iMSCs on soy-based scaffolds exhibited intracellular lipid droplet formation and positive Oil Red O staining (B). In contrast, iMSCs cultured on scaffolds under proliferation conditions showed no detectable Oil Red O staining (C). Representative images are shown for each condition. Scale bar = 100 µm.

Following adipogenic differentiation, ORO staining revealed accumulation of intracellular lipid droplets within iMSCs cultured and differentiated on soy-based scaffolds in shakerflask conditions (Figure 11B). In contrast, undifferentiated iMSC–scaffold cultures maintained under identical conditions showed no detectable ORO–positive staining (Figure 11C), indicating that lipid accumulation was differentiation-dependent rather than an artefact of scaffold association or dynamic culture.

Taken together, the observed biomass accumulation and preserved adipogenic differentiation highlight the suitability of this scaffold-based dynamic culture strategy for structured cultured tissue applications.

## Discussion

Cultivated meat manufacturing remains constrained by three linked challenges: (i) establishing renewable, wellcharacterized livestock cell sources that can be banked, expanded and differentiated to key target cell types reproducibly; (ii) achieving highdensity culture at scale while maintaining acceptable nutrient delivery and waste removal; and (iii) enabling edible structurization so that biomass can be formed into foodrelevant architectures efficiently and without costly scaffold removal steps. Recent technoeconomic syntheses emphasize that feasibility at industrial scale depends strongly on productivity and timeinculture, and that many published laboratory workflows still do not align with the assumptions used in scaleup models (Goodwin et al., 2024). In parallel, recent reviews of cultivated meat scaffolds stress that edible, nonanimal materials must satisfy both biological requirements (attachment, differentiation, mass transfer) and food constraints (safety, manufacturability, and ingredient compatibility) (Seibert et al., 2025; Wang et al., 2026). Finally, generating fat is increasingly recognized as essential for food relevance, because adipose tissue strongly affects organoleptic perception, and scaffold engineering can modulate differentiation quality and downstream foodrelevant attributes (Lee et al., 2024).

The present study demonstrates an integrated workflow that addresses each of these constraints by combining a renewable bovine ESC-derived iMSC platform with edible soy-based scaffolds and dynamic culture to achieve rapid biomass accumulation while retaining adipogenic differentiation capacity. The iMSC platform offers a reproducible starting point aligned with the need for stable, banked cell sources highlighted by techno-economic analysis (TEA) literature (Goodwin et al., 2024). At the materials level, soy-based scaffolds were selected because plant-protein scaffold systems are increasingly supported as food-compatible structuring substrates, and soy in particular has been reported to support strong adhesion and proliferation in stem-cell-relevant contexts (Kim et al., 2024). Consistent with that direction, our results show preferential bovine iMSC attachment, distribution, and early biomass accumulation on soy-based scaffolds, with pea-based scaffolds as an alternative regarding seeding, compared with lentil-based scaffolds, supporting soy as a practical edible scaffold material for short-term, high-density culture.

A key operational finding was that dynamic culture improved shortterm productivity relative to static controls. Under shakerflask conditions, cell-scaffold cultures exhibited increased glucose consumption and lactate production alongside increased biomass accumulation, consistent with the general observation that proliferative mammalian cells can display high glycolytic flux with lactate formation under common culture conditions (Hefzi et al., 2025). While mechanistic interpretation should remain conservative without pathwaylevel flux analysis, the observed increase in net biomass formation under dynamic culture conditions is consistent with improved effective mass transfer during the rapid expansion phase, which is precisely the type of processintensification lever highlighted as important for bridging bench workflows to manufacturingrelevant processes (Goodwin et al., 2024).

Beyond expansion, the study also demonstrates functional adipogenesis after scaffold-based culture, addressing a major gap in many structurization workflows where expansion and differentiation are not integrated in a food-compatible manner. Lipid accumulation was confirmed by staining following adipogenic induction. This aligns with recent work showing that scaffold engineering can be used to regulate differentiation quality and improve food-relevant attributes in cultured meat constructs (Lee et al., 2024). Our findings also sit alongside emerging evidence that dynamic 3D culture strategies can produce stable adipogenic building blocks suitable for downstream structuring approaches (Klatt et al., 2024). While composition of FA on scaffolds was not analyzed, evidence showing the nutritional similarity of differentiated adipogenic iMSCs to bovine tissue shows the potential of this cell source to provide lipids to improve the nutritional composition of a cell-scaffold food prototype. Importantly, our workflow links high-density scaffold culture directly to on-scaffold adipogenic differentiation, supporting a more integrated “culture-to-structured prototype” approach consistent with the direction suggested by recent scaffold and cultivated meat structurization reviews (Seibert et al., 2025; Wang et al., 2026).

In terms of edible scaffold choice and broader comparability, recent studies have shown soy-based edible scaffolds can support 3D muscle tissue formation and that scaffold microstructure can influence tissue yield and matrix deposition, reinforcing the broader potential of soy as a structuring substrate for cultivated meat rather than merely a carrier (Mariano et al., 2025). Complementary findings in cultivated meat bioprocess intensification (e.g., scalable operational strategies in serum-free contexts) further support the general principle that unit-operation optimization materially improves manufacturingrelevant outcomes (Bodiou et al., 2025). Taken together, these comparisons suggest that the key value of the present work is not simply demonstrating scaffold compatibility, but demonstrating a shortcycle, dynamic, edible scaffold workflow that enables cell expansion and differentiation for a production relevant output.

Several limitations should be considered when interpreting and extending these findings. Firstly, while adipogenesis and FA composition support functional lipid formation, broader nutritional equivalence would benefit from additional characterization beyond bulk FA profiles (e.g., lipid class distribution and stability under relevant processing conditions), as emphasized by recent organoleptic-focused cultivated meat studies (Lee et al., 2024). Secondly, because high lactate production can become inhibitory at larger scales, further work should define operating windows (agitation and feeding strategies, oxygen transfer) that sustain productivity while limiting accumulation of growth-limiting byproducts, consistent with broader mammalian cell bioprocess considerations (Hefzi et al., 2025). Similar shifts toward elevated glycolytic by-product formation under intensified or stress-associated culture conditions have been widely reported in mammalian cell systems, reinforcing that increased lactate production is a common feature of high-density culture rather than a system-specific metabolic mechanism (da Veiga Moreira et al., 2021). Thirdly, as scale increases, gradients and heterogeneity become more influential; this is repeatedly flagged as an open challenge in TEA literature and motivates future experiments in more controlled and scalable bioreactor formats (Goodwin et al., 2024).

Future experiments should therefore focus on: (i) systematic optimization of foodgrade media composition and feeding strategies to improve yield and reduce waste metabolite accumulation; (ii) fine-tuning of scaffold architecture (e.g., pore characteristics and surface properties) to further improve uniform cell distribution and maturation, consistent with scaffoldfocused research trends (Seibert et al., 2025; Wang et al., 2026); and (iii) expanding foodrelevant characterization toward texture evolution and sensorylinked chemistry, building on prior evidence that differentiation quality and matrix cues can influence organoleptic outcomes (Lee et al., 2024). Collectively, the results support an integrated strategy in which renewable bovine iMSC banks, edible soybased scaffolds, and dynamic culture conditions combine to accelerate structured biomass generation in a manner aligned with current scaleup and structurization priorities in the cultivated meat field.

## Data Availability Statement

The data supporting the findings of this study are available within the article and its supplementary materials. Sequence data that can be downloaded from Roslin Technologies servers on request from the corresponding author. Proprietary media and scaffolds are available from Roslin Technologies and sallea AG.

## Ethics Statement

The bovine embryonic stem cell lines used in this study were provided by Genus plc and was generated, banked, and supplied under the appropriate regulatory approvals, licenses, and ethical oversight in accordance with relevant national and international guidelines. All work described in this manuscript used previously established cell lines; no new animal-derived material was generated, and no live animal procedures were performed as part of this study.

## Author Contributions

Madeleine Carter, Tim Spitters, and Deepika Rajesh contributed to the conceptualization of the study. Methodology was developed by Madeleine Carter and Tim Spitters. Investigation was carried out by Sarah Ho on adipogenesis, Sophia Webb on RNAseq analysis, and Niamh Hyland for the provision of cells and tissue culture expertise. Formal oversight was performed by Patrick Joseph Mee and Deepika Rajesh. Resources and data relating to scaffolds were provided by Simona Fehlmann. All authors contributed to construction, reviewing and editing the manuscript. All authors have read and approved the final manuscript.

## Acknowledgments

We thank colleagues at Roslin Technologies and sallea AG for discussions and technical assistance. We are grateful to Multus for supply of food-grade basal media. Additionally, we thank the LABMaiTE team for the insightful discussions on data analysis and interpretation. We thank CADFEM for their input on the development of the CFD models. BioRender.com was used to generate the schematics in Figures 1, 2, 8, 10 and 11.

---

**Supplementary Figure S1.**
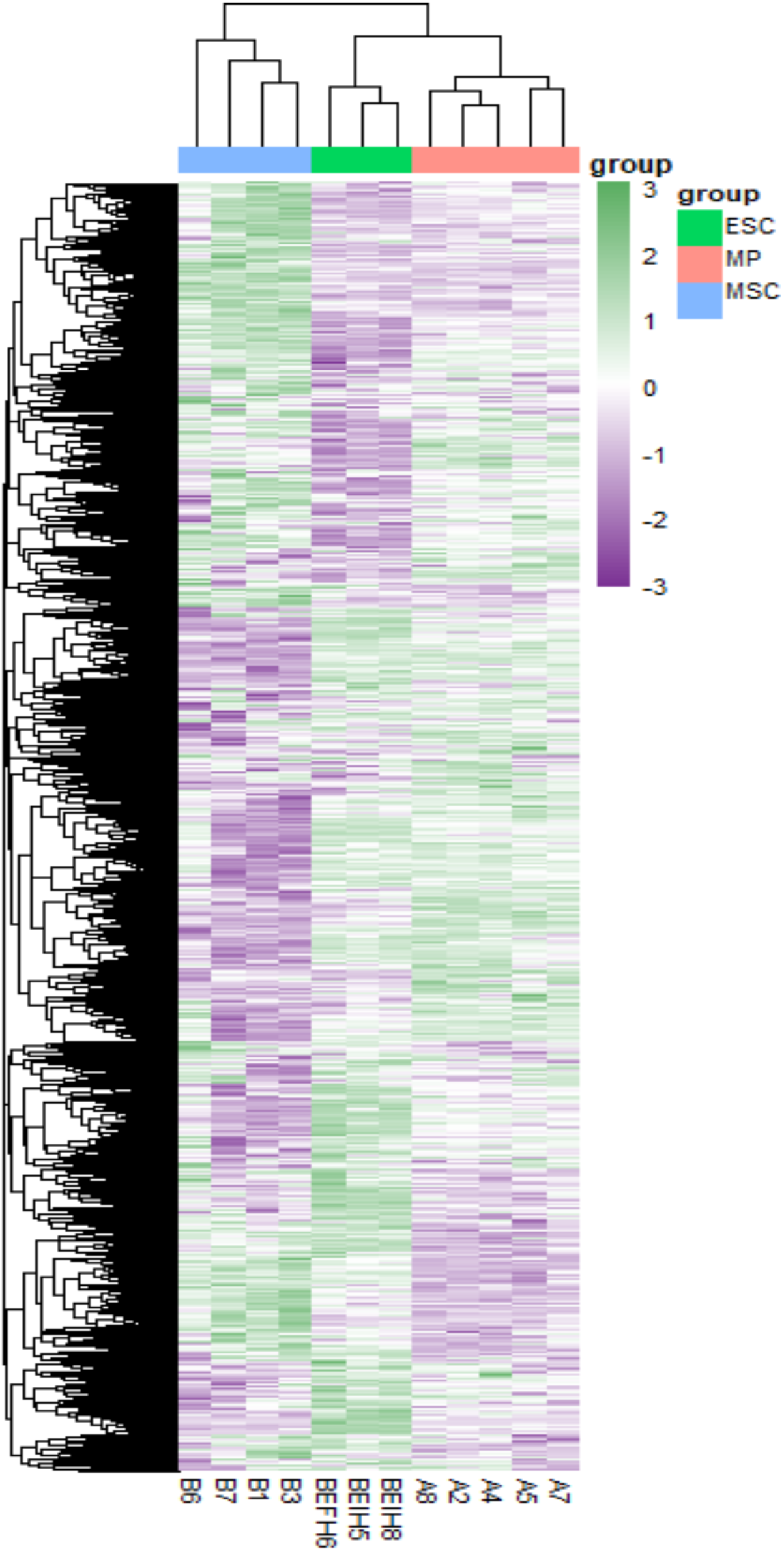
Hierarchical clustering of transcriptomic profiles from bovine embryonic stem cell–derived differentiation stages. Unsupervised hierarchical clustering of RNA-sequencing data from bovine embryonic stem cells (ESCs), mesodermal progenitors (MPs), and induced mesenchymal stem cells (iMSCs). Expression values are shown as log₂ counts per million (log₂CPM), with gene-wise normalisation to z-scores. Rows (genes) and columns (samples) were clustered independently to visualize similarities in transcript abundance across differentiation stages, with colour indicating relative expression levels.

**Supplementary Figure S2.**
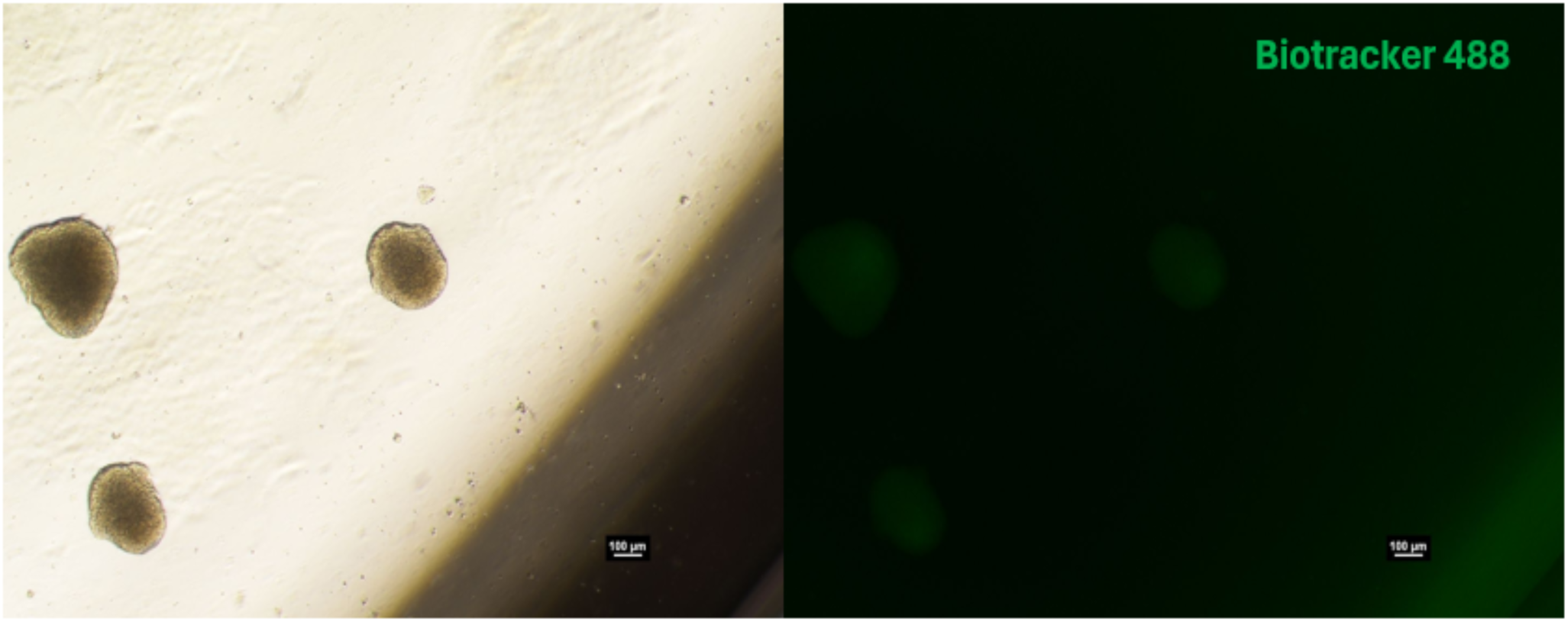
Lipid staining of bovine induced mesenchymal stem cells in suspension culture. Bovine induced mesenchymal stem cells (iMSCs) cultured under suspension conditions were stained with a fluorescent lipid tracer to assess intracellular lipid accumulation. Representative fluorescence images show iMSCs labelled with BioTracker™ 488 lipid dye, with no detectable signal corresponding to lipid droplet formation. Scale bar = 100 µm.

**Supplementary Figure S3.**
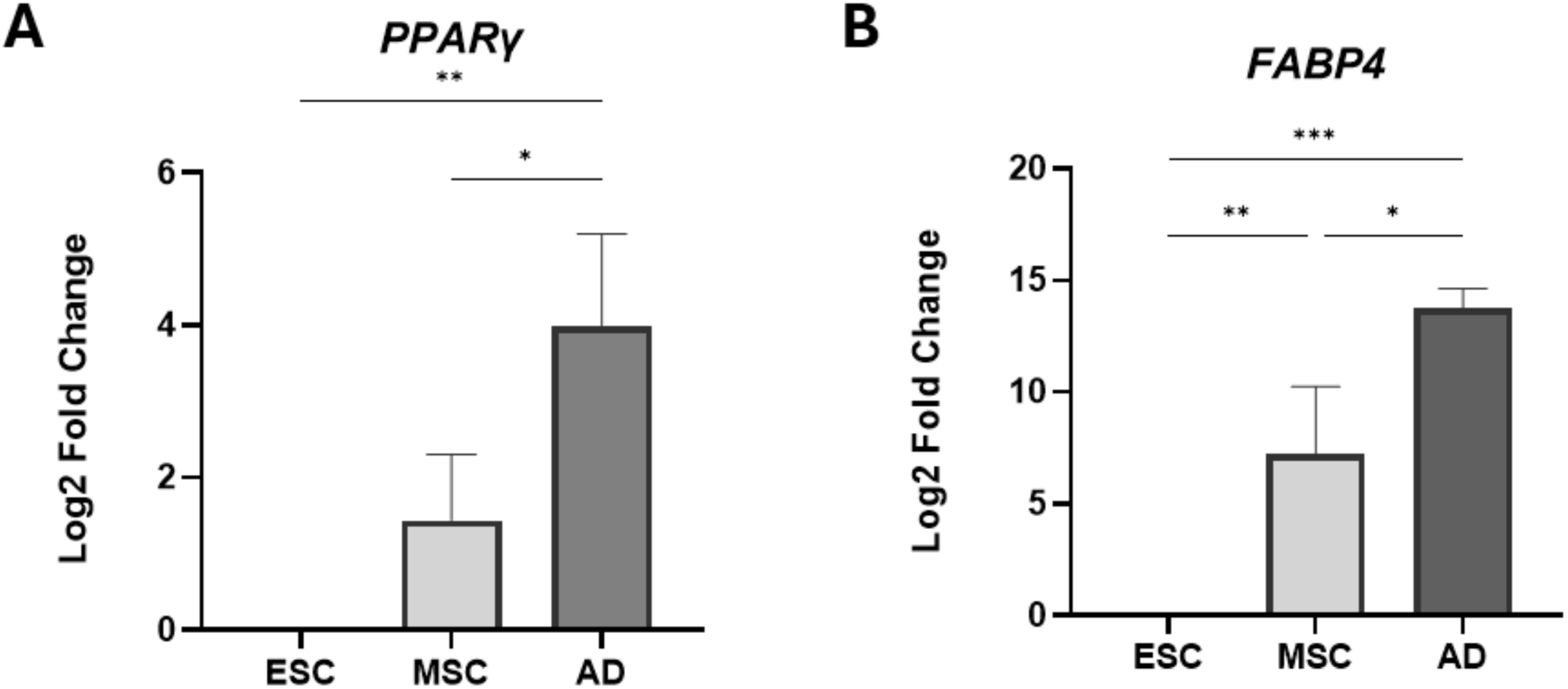
Adipogenic marker expression in differentiated three-dimensional bovine mesenchymal stem cell cultures. Expression of adipogenic marker genes was assessed in three-dimensional adipocyte cultures derived from bovine induced mesenchymal stem cells (iMSCs). Relative expression of *PPARγ* is increased compared to undifferentiated embryonic stem cells (p<0.0033) and iMSCs (p<0.0265) (n=3), (A), and *FABP4* is significantly increased compared to undifferentiated ESC (p<0.0002) and iMSCs (p<0.0107) (n=3) (B).

## References

Ben-Arye, T., Shandalov, Y., Ben-Shaul, S., Landau, S., Zagury, Y., Ianovici, I., Lavon, N., & Levenberg, S. (2020). Textured soy protein scaffolds enable the generation of three-dimensional bovine skeletal muscle tissue for cell-based meat. Nature Food, 1, 210–220. 10.1038/s43016-020-0046-5

Bodiou, V., Cristini, N., De Cristofaro, L., Pareek, T., Rajagopal, V., Verrougstraete, L., Heinrich, J. M., Post, M. J., & Moutsatsou, P. (2025). Process intensification of cultivated meat production through microcarrier addition strategy optimisation. Scientific Reports, 15, 14080. 10.1038/s41598-025-97813-7

Bosnakovski, D., Mizuno, M., Kim, G., Takagi, S., Okumura, M., & Fujinaga, T. (2005). Isolation and multilineage differentiation of bovine bone marrow mesenchymal stem cells. Cell and Tissue Research, 319(2), 243–253. 10.1007/s00441-004-1012-5

Chen, Y., Chen, L., Lun, A. T. L., Baldoni, P. L., & Smyth, G. K. (2025). edgeR v4: Powerful differential analysis of sequencing data with expanded functionality and improved support for small counts and larger datasets. Nucleic Acids Research, 53(2), gkaf018. 10.1093/nar/gkaf018

da Veiga Moreira, J., De Staercke, L., Martínez-Basilio, C., Gauthier-Thibodeau, S., Montégut, L., Schwartz, L., & Jolicoeur, M. (2021). Hyperosmolarity triggers the Warburg effect in Chinese hamster ovary cells and reveals a reduced mitochondrial horsepower. Metabolites, 11(6), 344. 10.3390/metabo11060344

De la Garza-Rodea, A. S., van der Velde-van Dijke, I., Boersma, H., Gonçalves, M. A. F. V., van Bekkum, D. W., de Vries, A. A. F., & Knaan-Shanzer, S. (2012). Myogenic properties of human mesenchymal stem cells derived from three different sources. Cell Transplantation, 21(1), 153–173. 10.3727/096368911X580554

Dobin, A., Davis, C. A., Schlesinger, F., Drenkow, J., Zaleski, C., Jha, S., Batut, P., Chaisson, M., & Gingeras, T. R. (2013). STAR: Ultrafast universal RNA-seq aligner. Bioinformatics, 29(1), 15–21. 10.1093/bioinformatics/bts635

Dohmen, R. G. J., Hubalek, S., Melke, J., Messmer, T., Cantoni, F., Mei, A., Hueber, R., Mitic, R., Remmers, D., Moutsatsou, P., Post, M. J., Jackisch, L., & Flack, J. E. (2022). Muscle-derived fibroadipogenic progenitor cells for production of cultured bovine adipose tissue. npj Science of Food, 6, Article 1. 10.1038/s41538-021-00122-2

Ebner-Peking, P., Krisch, L., Wolf, M., Hochmann, S., Hoog, A., Vári, B., Muigg, K., Poupardin, R., Scharler, C., Schmidhuber, S., Russe, E., Stachelscheid, H., Schneeberger, A., Schallmoser, K., & Strunk, D. (2021). Self-assembly of differentiated progenitor cells facilitates spheroid human skin organoid formation and planar skin regeneration. Theranostics, 11(17), 8430–8447. 10.7150/thno.59661

Fish, K. D., Rubio, N. R., Stout, A. J., Yuen, J. S. K., & Kaplan, D. L. (2020). Prospects and challenges for cell-cultured fat as a novel food ingredient. Trends in Food Science & Technology, 98, 53–67. 10.1016/j.tifs.2020.02.005

Fraeye, I., Kratka, M., Vandenburgh, H., & Thorrez, L. (2020). Sensorial and nutritional aspects of cultured meat in comparison to traditional meat: Much to be inferred. Frontiers in Nutrition, 7, 35. 10.3389/fnut.2020.00035

Goodwin, C. M., Aimutis, W. R., & Shirwaiker, R. A. (2024). A scoping review of cultivated meat techno-economic analyses to inform future research directions for scaled-up manufacturing. Nature Food, 5(11), 901–910. 10.1038/s43016-024-01061-3

Hanga, M. P., Fehlmann, S., Kleger, N., Vieri, M. L., Callue, T., Millbank, A., & Theodosiou, E. (2025). Edible scaffolds for cultivated meat production. In Advances in Biochemical Engineering/Biotechnology. Springer Nature. 10.1007/10_2025_291

Hill, A. B. T., Bressan, F. F., Murphy, B. D., & Garcia, J. M. (2019). Applications of mesenchymal stem cell technology in bovine species. Stem Cell Research & Therapy, 10(1), 44. 10.1186/s13287-019-1145-9

Hefzi, H., Martínez-Monge, I., Marin de Mas, I., Cowie, N. L., Gomez Toledo, A., Noh, S. M., et al. (2025). Multiplex genome editing eliminates lactate production without impact on growth rate in mammalian cells. Nature Metabolism, 7, 212–227. 10.1038/s42255-024-01193-7

Kim, D. H., Han, S. G., Lim, S. J., Hong, S. J., Kwon, H. C., Jung, H. S., & Han, S. G. (2024). Comparison of soy and pea protein for cultured meat scaffolds: Evaluating gelation, physical properties, and cell adhesion. Food Science of Animal Resources, 44(5), 1108–1125. 10.5851/kosfa.2024.e46

Klatt, A., Wollschlaeger, J. O., Albrecht, F. B., Rühle, S., Holzwarth, L. B., Hrenn, H., Melzer, T., Heine, S., & Kluger, P. J. (2024). Dynamically cultured, differentiated bovine adipose-derived stem cell spheroids as building blocks for biofabricating cultured fat. Nature Communications, 15, 9107. 10.1038/s41467-024-53486-w

Lee, M., Park, S., Choi, B., Choi, W., Lee, H., Lee, J. M., et al. (2024). Cultured meat with enriched organoleptic properties by regulating cell differentiation. Nature Communications, 15, 77. 10.1038/s41467-023-44359-9

Lew, E. T., Yuen, J. S. K., Zhang, K. L., Fuller, K., Frost, S. C., & Kaplan, D. L. (2024). Chemical and sensory analyses of cultivated pork fat tissue as a flavor enhancer for meat alternatives. Scientific Reports, 14(1), 17643. 10.1038/s41598-024-68247-4

Lewis-Israeli, Y. R., Wasserman, A. H., Gabalski, M. A., Volmert, B. D., Ming, Y., Ball, K. A., et al. (2021). Self-assembling human heart organoids for the modeling of cardiac development and congenital heart disease. Nature Communications, 12(1), 5142. 10.1038/s41467-021-25329-5

Li, C.-H., Yang, I.-H., Ke, C.-J., Chi, C.-Y., Matahum, J., Kuan, C.-Y., Celikkin, N., Swieszkowski, W., & Lin, F.-H. (2022). The production of fat-containing cultured meat by stacking aligned muscle layers and adipose layers formed from gelatin–soymilk scaffold. Frontiers in Bioengineering and Biotechnology, 10, 875069. 10.3389/fbioe.2022.875069

Liao, Y., Smyth, G. K., & Shi, W. (2014). featureCounts: An efficient general purpose program for assigning sequence reads to genomic features. Bioinformatics, 30(7), 923–930. 10.1093/bioinformatics/btt656

Mariano, E., Lee, D. Y., Choi, Y., Park, J., Han, D., Kim, J. S., Park, J. W., Namkung, S., Joo, S.-T., Choi, I., & Hur, S. J. (2025). Crusting-fabricated soy protein-based scaffolds yield three-dimensional muscle tissues for cultured chicken meat production. Journal of Food Science, 90(4), e70139. 10.1111/1750-3841.70139

Martin, M. (2011). Cutadapt removes adapter sequences from high-throughput sequencing reads. EMBnet.journal, 17(1), 10–12. 10.14806/ej.17.1.200

Moslemy, N., Sharifi, E., Asadi-Eydivand, M., & Abolfathi, N. (2023). Review in edible materials for sustainable cultured meat: Scaffolds and microcarriers production. International Journal of Food Science & Technology, 58(12), 6182–6191. 10.1111/ijfs.16703

Seibert, G. A., Feddern, V., Bastos, A. P. A., Kumar, A., & Verruck, S. (2025). Trends in non-animal scaffolds for cultured meat structuration. NPJ Science of Food, 9, Article 208. 10.1038/s41538-025-00429-4

Schmittgen, T. D., & Livak, K. J. (2008). Analyzing real-time PCR data by the comparative CT method. Nature Protocols, 3(6), 1101–1108. 10.1038/nprot.2008.73

Wang, Y., Zou, L., Liu, W., & Chen, X. (2023). An overview of recent progress in engineering three-dimensional scaffolds for cultured meat production. Foods, 12(13), 2614. 10.3390/foods12132614

Ward, S. R., Tomiya, A., Regev, G. J., Thacker, B. E., Benzl, R. C., Kim, C. W., & Lieber, R. L. (2009). Passive mechanical properties of the lumbar multifidus muscle support its role as a stabilizer. Journal of Biomechanics, 42(10), 1384–1389. 10.1016/j.jbiomech.2008.09.042

Yao, Y., Yuen, J. S. K., Jr., Sylvia, R., Fennelly, C., Cera, L., Zhang, K. L., Li, C., & Kaplan, D. L. (2024). Cultivated meat from aligned muscle layers and adipose layers formed from glutenin films. ACS Biomaterials Science & Engineering, 10(2), 814–824. 10.1021/acsbiomaterials.3c01500

